# Machine learning-guided olivetolic acid cyclase engineering enables tailored cannabinoid biosynthesis in yeast

**DOI:** 10.64898/2026.06.12.731972

**Authors:** Nathaniel Blalock, Joseph W LaMattina, Emma Monge, Raymond Tran, Audrey E. Louie, Jun Urano, Spiros Kambourakis, Russell S Komor, Philip A Romero

## Abstract

Cannabinoids comprise a diverse class of bioactive natural products with important therapeutic potential, but efficient microbial production remains limited by pathway bottlenecks and challenges in engineering key biosynthetic enzymes. Here, we develop a machine learning-guided approach to engineer olivetolic acid cyclase (OAC), a critical control point in cannabinoid biosynthesis that governs both pathway flux and product selectivity. We first generated sequence-function data from 152 CsOAC variants spanning homolog screening, recombination, and mutagenesis libraries. Using these measurements, we trained multi-task models to predict pathway-level production of olivetolic acid (OA), divarinic acid (DVA), and competing byproducts, together with a variational autoencoder that captured evolutionary constraints across the broader enzyme family. Across three rounds of iterative design and testing, this approach identified CsOAC variants that substantially increased production and selectivity of both OA and DVA. When introduced into engineered *Yarrowia lipolytica* strains, these variants enabled production of tetrahydrocannabinolic acid (THCA) and the minor cannabinoid tetrahydrocannabivarinic acid (THCVA) at titers exceeding previous yeast systems. Analysis of top-performing variants revealed mutations influencing substrate selectivity and catalytic performance, providing insight into the determinants of CsOAC function. More broadly, this work demonstrates how machine learning-guided enzyme engineering can improve pathway performance and expand access to major and minor cannabinoids through microbial biosynthesis.

## Introduction

Cannabinoids comprise a structurally diverse class of natural products with broad therapeutic potential, including clinically approved compounds such as Δ^9^-tetrahydrocannabinol (THC) and cannabidiol (CBD) for the treatment of pain, epilepsy, and other neurological disorders^1–6^. Beyond these major compounds, numerous minor cannabinoids and analogues exhibit distinct pharmacological properties, highlighting the importance of accessing expanded cannabinoid chemical space^7–10^. Microbial biosynthesis has emerged as a powerful platform to produce cannabinoids in a scalable and controlled manner^11,12^, with heterologous pathway reconstitution demonstrated in hosts such as *Escherichia coli*^13–15^, *Saccharomyces cerevisiae*^16,17^, and *Yarrowia lipolytica*^18,19^. These systems also enable the production of non-natural cannabinoid analogues by introducing alternative precursors. However, a key and underexploited opportunity lies in engineering pathway enzyme specificity to directly control product structure, enabling selective production of major and minor cannabinoids as well as potentially new-to-nature analogues that are difficult to access through plant extraction or chemical synthesis.

A key constraint in cannabinoid biosynthesis is olivetolic acid cyclase from *Cannabis sativa* (CsOAC), which catalyzes cyclization of a reactive tetraketide intermediate to form olivetolic acid, the central precursor to all major cannabinoids. Early biochemical studies established that CsOAC is required for olivetolic acid formation^20^, and subsequent pathway engineering efforts showed that increasing CsOAC expression alone can improve product titers^16^, underscoring its control over pathway flux. Mechanistically, CsOAC competes with non-enzymatic hydrolysis of the unstable tetraketide intermediate and lactonization, leading to the formation of derailment products that reduce pathway efficiency and may inhibit downstream steps. In addition to controlling flux, CsOAC influences product distribution through substrate^16,21^ selectivity, particularly for non-native fatty acid–derived intermediates^16^. Despite its importance, CsOAC has proven difficult to engineer. Functional homologs are rare, limiting opportunities for discovery-based approaches, and rational design is challenged by the enzyme’s role within a multi-step pathway involving unstable intermediates and competing reactions. Prior work has investigated targeted substitutions to CsOAC, but these efforts yielded only limited improvements^22^. These features suggest that improving CsOAC requires approaches that can learn sequence–function relationships directly from pathway-level measurements rather than relying on isolated biochemical assays.

Here, we develop a machine learning–guided approach to engineer CsOAC for improved activity and substrate selectivity within the cannabinoid biosynthetic pathway. To enable this, we first establish a scalable microbial platform for cannabinoid biosynthesis in *Y. lipolytica* through extensive pathway reconstruction and host engineering, enabling production of both native and minor cannabinoids from fatty acid–derived precursors. Using this system, we identify CsOAC as a key control point governing pathway flux and product distribution. We then engineer CsOAC by combining experimentally derived sequence–function data with multi-objective models that jointly optimize product formation and pathway selectivity, together with an evolutionary prior to constrain designs to functional regions of sequence space. Across iterative rounds of design and testing, this approach yields CsOAC variants that substantially improve production of desired cannabinoid precursors and enhance downstream cannabinoid biosynthesis. More broadly, this work demonstrates how machine learning–guided enzyme engineering can overcome key bottlenecks in complex biosynthetic pathways and expand access to new-to-nature small molecules.

## Results

### Production of cannabinoids in oleaginous yeast

*Cannabis sativa* uniquely produces more than 100 diverse cannabinoids^23,24^, but plant-based production is slow, difficult to scale, and largely constrained to native biosynthetic pathways and naturally occurring structures^6,11,16^. Microbial biosynthesis offers a scalable and controllable alternative, with the potential to access non-natural cannabinoid analogues^11^. Prior work has demonstrated transplantation of cannabinoid pathways into hosts such as *E. coli*^13–15^, *S. cerevisiae*^16,17^, and *Y. lipolytica*^18,19^, but these systems have generally been limited to precursor production and have achieved relatively low titers. Despite these limitations, *Y. lipolytica* remains an attractive host for cannabinoid biosynthesis because of its high flux through acetyl-CoA and malonyl-CoA, efficient lipid metabolism, and ability to sequester hydrophobic metabolites such as cannabinoids in lipid droplets. In addition, its eukaryotic machinery supports proper folding and post-translational modification of plant-derived enzymes. Here, we leverage these features to engineer improved cannabinoid production strains.

In *C. sativa*, cannabinoid biosynthesis proceeds through a well-defined pathway that converts fatty acid–derived precursors into bioactive cannabinoids6/12/26 3:04:00 PM^16,25^. The pathway is initiated from hexanoic acid, which is activated to hexanoyl-CoA by an acyl-activating enzyme (CsAAE1)^16^. A type III polyketide synthase, tetraketide synthase (CsPKS)^26^, then catalyzes sequential condensations of hexanoyl-CoA with three malonyl-CoA units to generate a reactive linear tetraketide intermediate (3,5,7-trioxododecanoyl-CoA), while the non-enzymatic hydrolysis of earlier polyketide intermediates can divert flux to derailment products such as pentyl diacetic acid lactone (PDAL). The tetraketide intermediate is subsequently cyclized by olivetolic acid cyclase (CsOAC)^20^ to form olivetolic acid (OA), a key pathway intermediate whose efficient conversion is critical to prevent flux loss to the competing side reaction to form olivetol (OL). OA is prenylated by an aromatic prenyltransferase (CsAPT)^16^ using geranyl-pyrophosphate (GPP) to produce the central intermediate cannabigerolic acid (CBGA). From CBGA, downstream oxidocyclase enzymes determine final product identity: tetrahydrocannabinolic acid synthase (CsTHCAS)^27,28^ and cannabidiolic acid synthase (CsCBDAS) convert CBGA into Δ9-tetrahydrocannabinolic acid (THCA) and cannabidiolic acid (CBDA), respectively^16^. These acidic cannabinoids can subsequently undergo non-enzymatic decarboxylation to yield the neutral cannabinoids tetrahydrocannabinol (THC) and cannabidiol (CBD).

We reconstructed the core *C. sativa* pathway in *Y. lipolytica* to enable cannabinoid biosynthesis in a microbial host with stable genomic integration of key enzymes spanning precursor activation, polyketide formation, cyclization, and downstream oxidation (Figure 1, Supplementary Table 1-2). We introduced an engineered tetraketide synthase (CsPKS1.1) containing a mutation that improved OA and divarinic acid (DVA) titers (Supplementary Figure 1), CsOAC, and a tetrahydrocannabinolic acid synthase modified for expression in *Y. lipolytica* (CsTHCAS*)^25^ to establish the core route from fatty acid–derived intermediates to THCA. To enable efficient activation of medium-chain fatty acids, we incorporated an engineered hexanoyl-CoA synthetase derived from *Arabidopsis thaliana* (AtHCS2)^29^. We replaced the native *C. sativa* aromatic prenyltransferase with an engineered variant from *Streptomyces longwoodensis* (SlAPT73.248), selected for its soluble expression, improved selectivity for geranyl pyrophosphate, and broader tolerance toward non-native cannabinoid precursors^30^. Together, this engineered pathway enables conversion of exogenously supplied fatty acids into the central cannabinoid intermediate CBGA and onward to THCA within a heterologous yeast chassis, establishing a functional end-to-end biosynthetic route in *Y. lipolytica*.

**Figure 1:**
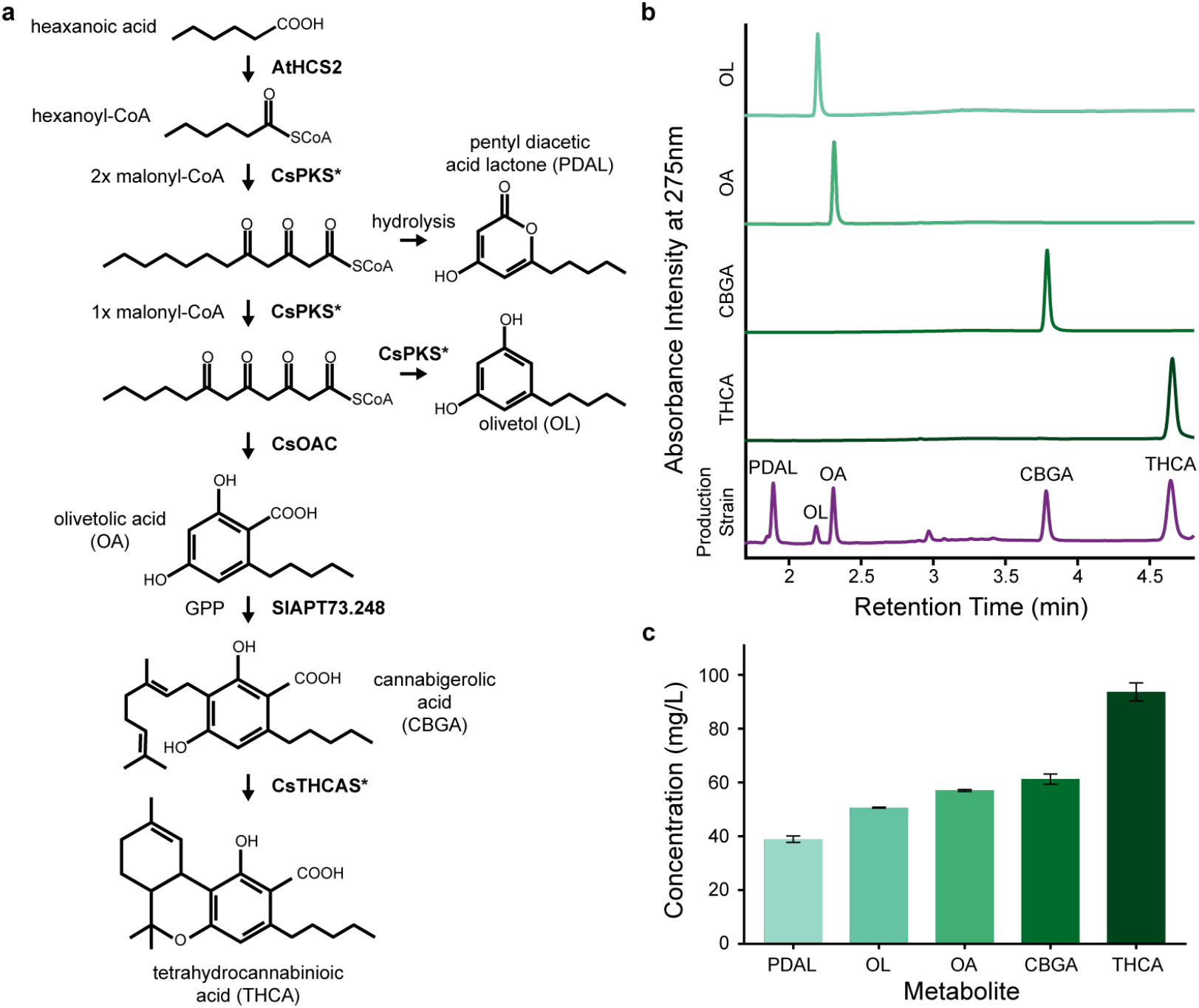
Production of cannabinoids in oleaginous yeast. **a**, Expression of AtHCS2 enables production of hexanoyl-CoA or butyryl-CoA after from hexanoic acid or butyric acid feeds. Expression of CsPKS* and CsOAC enables production of OA or DVA from hexanoyl-CoA or butyryl-CoA. After two malonyl-CoA condensations, the reactive linear triketide intermediate can undergo hydrolysis to produce the undesirable byproduct pentyl diacetic acid lactone (PDAL). After three malonyl-CoA condensations, the reactive linear tetraketide product undergoes competition between decarboxylative aldol-condensation to produce the undesirable byproduct OL and cyclization by CsOAC to produce OA. Production of byproducts PDAL and OL reduce pathway flux and can reduce growth^36^. Expression of SlAPT73.248 enables production of CBGA and cannabigerovarinic acid (CBGVA) via the prenylation of OA or DVA. The expression of CsTHCAS* enables production of THCA or tetrahydrocannabivarinic acid (THCVA). Engineered variants of CsPKS and CsTHCAS are indicated with an asterisk (Supplementary Table 1-2). GPP production is described in Supplementary Figure 2 and Supplementary Table 1-3. **b**, LC-DAD chromatograms at 275nm for OL standard, OA standard, CBGA standard, THCA standard, and a *Y. lipolytica* production strain containing the full cannabinoid pathway with peaks confirming the production of PDAL, OL, OA, CBGA, and THCA. There was no commercially available standard for PDAL. **c**, Concentrations of PDAL, OL, OA, CBGA, and THCA in the production strain, calculated by integrating the LC-DAD peaks from b and interpolating against full calibration curves. Bars show mean concentration in mg/L; error bars indicate standard deviation from three biological replicates (n = 3).

To support high-level cannabinoid production, we extensively reprogrammed the metabolism of *Y. lipolytica* to increase precursor supply, optimize pathway flux, and enable efficient expression of heterologous enzymes (Supplementary Figure 2, Supplementary Table 1-3). We enhanced cytosolic acetyl-CoA availability through overexpression of native acetyl-CoA synthetase (YlACS1)^31^ and increased flux through the mevalonate pathway via upregulation of HMG-CoA reductase (YlHMGR)^32,33^, thereby supporting production of the isoprenoid precursor GPP^31^. To further bias flux toward GPP, we introduced two engineered variants of the native *Y. lipolytica* GPP/FPP synthase YlErg20: YlErg20.F88W.N119W that has been shown to enhance linalool production in *Y. lipolytica*^34^ and YlErg20.A28, our engineered variant that favors GPP accumulation while maintaining pathway throughput^35^. In parallel, we minimized loss of key intermediates by disrupting β-oxidation genes (acd1, pox3, and pox5) to prevent degradation of medium-chain acyl-CoAs and by perturbing branched-chain fatty acid metabolism (ivd1 and bckd) to redirect flux toward C4-CoA species and reduce formation of off-pathway byproducts^31^. Finally, to support functional expression of plant-derived enzymes, we enhanced endoplasmic reticulum folding capacity through overexpression of the chaperone YlCNE1^25^. Together, these modifications establish a metabolically optimized chassis that integrates precursor supply, pathway flux, and protein expression to enable efficient cannabinoid biosynthesis.

We next evaluated cannabinoid production in the fully engineered *Y. lipolytica* strain using fed-batch fermentations supplemented with either hexanoic acid or butyric acid to drive C6- and C4-derived pathways, respectively. Targeted metabolomic analysis confirmed that the reconstructed pathway was functional end-to-end, with production of cannabinoid intermediates and final products including THCA (Figure 1b). Despite this successful pathway reconstitution, substantial accumulation of upstream intermediates and off-pathway byproducts was observed, particularly under C4 feeding conditions. These byproducts are consistent with hydrolytic shunting of the reactive tetraketide intermediate, which is known to be chemically unstable. We therefore hypothesized that insufficient catalytic turnover by olivetolic acid cyclase (OAC) allows this intermediate to escape productive cyclization, leading to loss of flux and reduced pathway efficiency. This observation identifies CsOAC as a key bottleneck in cannabinoid biosynthesis and motivates its targeted engineering to improve both pathway flux and product selectivity.

### Learning CsOAC sequence-function relationships

We initially applied several protein engineering strategies to engineer CsOAC activity and substrate selectivity toward OA and its less preferred analogue DVA, which is derived from a C4 precursor rather than the native C6 precursor. Although CsOAC is named for its native olivetolic acid product, the enzyme can also cyclize the corresponding C4-derived tetraketide intermediate to produce DVA. For clarity, we refer to the enzyme and its variants as CsOAC throughout this study. We screened 40 CsOAC homologs for OA and DVA production, but none produced either metabolite above the analytical limit of detection, consistent with previous observations that OA/DVA-producing activity is not broadly distributed among known homologs^37,38^. We next created recombination variants by swapping secondary-structure elements from selected homologs into the *C. sativa* CsOAC scaffold to perturb active site pocket size and substrate selectivity. These donor sequences included a *C. sativa* homolog that produced low levels of OA/DVA *in vitro* (but not in *Y. lipolytica*) and a *Rhododendron dauricum* homolog that natively produces 1-carbon tail analogue orsellinic acid, but neither enhanced OA/DVA production (Figure 2a-b). We compared the CsOAC crystal structure with predicted donor homolog structures obtained with AlphaFold3^39,40^. Homologs had smaller predicted substrate pocket volumes than CsOAC, indicating scaffolds from the related donor homologs were not well suited for OA/DVA production^41^ (Supplementary Figure 3). Given the lack of success sampling the diversity of natural sequences related to CsOAC, we performed single site-saturation mutagenesis of residues in the CsOAC active site, but we observed limited improvements in OA/DVA production (Figure 2a-b). Together, these efforts produced an initial dataset of 152 CsOAC sequences spanning a wide range of sequence perturbations, providing a foundation for learning CsOAC sequence-function relationships within the cannabinoid biosynthetic pathway (Supplementary Figure 4, Supplementary Table 4).

**Figure 2:**
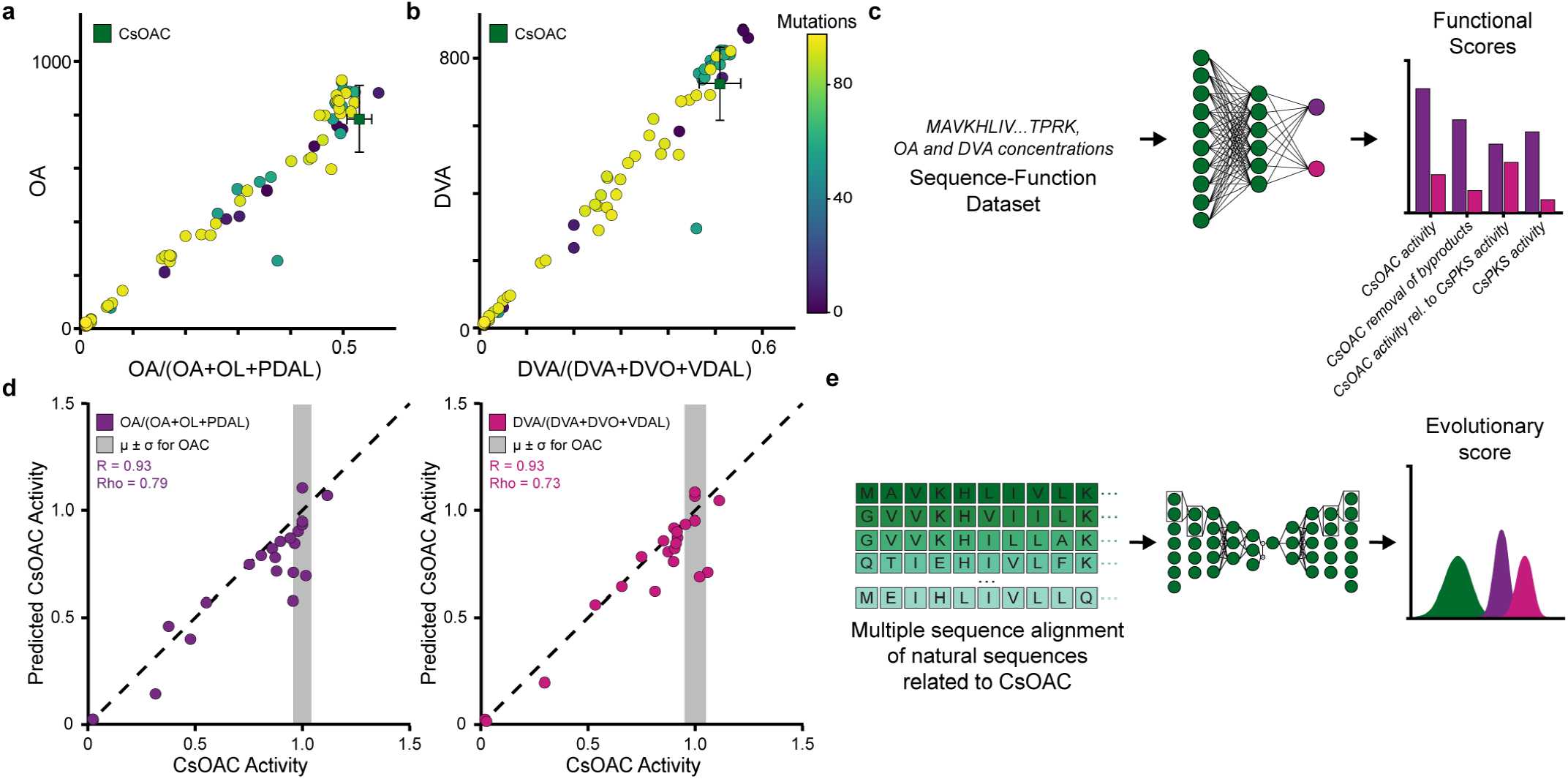
Learning CsOAC sequence-function relationships. **a-b**, Experimental characterization of initial CsOAC sequence variants for used for model training. OA titers versus pathway selectivity OA/(OA+OL+PDAL) **a**, and DVA titers versus DVA/(DVA+DVO+VDAL). Key: divarinol (DVO), varinic diacetic acid lactone (VDAL). **b**, illustrates functional diversity across sequences. **c**, Multi-task neural network predict OA (purple) and DVA (pink) production learned from experimentally characterized variants. **d**, Multi-task neural network predictions demonstrate strong rank-order agreement to unseen variants sequences withheld from training for predicted enzyme activity for OA/(OA+OL+PDAL) (purple) and DVA/(DVA+DVO+VDAL) (pink) versus experimental enzyme activity. **e**, Variational autoencoder (VAE) trained on natural homologs learns evolutionary constraints defining plausible enzyme sequence space, providing an evolutionary prior complementary to functional prediction.

We trained an ensemble of multi-task neural networks to learn CsOAC sequence–function relationships from this initial dataset by jointly predicting OA, DVA and their corresponding C6- and C4-derived byproducts (Figure 2c, Supplementary Figure 5, Supplementary Table 5). A multi-task modeling approach was essential for two reasons. First, it leverages the shared structure across cannabinoid pathway outputs, capturing tradeoffs between product and byproduct formation, enabling downstream prediction of pathway relevant ratios rather than modeling each metabolite in isolation. Second, a multi-task modeling approach naturally accommodates incomplete measurements, allowing us to train on all available data without discarding partially characterized sequences, since the sequence variants were not consistently assayed for both OA and DVA production (Supplementary Figure 6). The ensemble achieved strong rank-order agreement with experiments with a Spearman’s rank correlation coefficient of 0.79 for OA selectivity and 0.73 for DVA selectivity for withheld sequences, providing a functional predictor to guide CsOAC evolution (Figure 2d).

While the supervised model learned sequence-function relationships from experimental measurements, we sought a complementary model that captured evolutionary constraints across the broader enzyme family. We therefore trained a variational autoencoder (VAE) on a multiple sequence alignment of natural sequences related to CsOAC to provide an evolutionary prior for sequence design (Figure 2e, Supplementary Table 6, Supplementary Figure 7). The VAE evolutionary score quantifies how well candidate sequences conform to statistical constraints learned from natural homologs. Notably, CsOAC itself lies in the 0.041 quantile of MSA evolutionary scores, consistent with experimental observations that OA production is not broadly conserved among homologous polyketide cyclase sequences (Supplementary Figure 8a). Introducing mutations to CsOAC rapidly decreased evolutionary scores, with only 13.7% of single substitutions improving upon CsOAC and this fraction decaying to zero with the introduction of 6 mutations, indicating a highly constrained and epistatic sequence–function landscape (Supplementary Figure 8b-c). This trend mirrors our initial sequence libraries, where the accumulation of rational mutations reduced CsOAC activity. To assess whether the evolutionary prior captured mechanistically meaningful constraints, we examined mutations at F24 and V59, residues lining the substrate-binding pocket previously implicated in substrate binding affinity^37^. The VAE correctly identified V59I as favorable and V59A as deleterious^37,42^ (Supplementary Figure 8d). Similarly, F24I was predicted to be tolerated and has been experimentally associated with modest improvements in analogue production^42^ (Supplementary Figure 8d). These results demonstrate the VAE captures family-specific evolutionary constraints relevant to CsOAC enzyme stability and substrate binding that can complement the functional predictor to guide CsOAC evolution.

### Machine learning-guided evolution of CsOAC activity and specificity

Having learned both functional and evolutionary constraints on CsOAC sequence space, we next sought to determine whether these models could guide the design of improved enzyme variants. Because we wished to optimize both predicted function and evolutionary plausibility simultaneously, we framed sequence design as a multi-objective optimization problem. We performed three rounds of machine learning-guided CsOAC evolution. In each round, we retrained the multi-task neural network with all available experimental data (Supplementary Figure 9-10) and generated sequence designs along the Pareto front between functional and evolutionary model scores (Figure 3a, Supplementary Figure 11-12, Supplementary Table 7-8). Exploring the tradeoff between these complementary objectives enabled broader exploration of the CsOAC sequence-function landscape while maintaining evolutionary plausibility (Figure 3b-c). Selected designs were then experimentally characterized and the resulting measurements used to update the models for the subsequent round.

**Figure 3:**
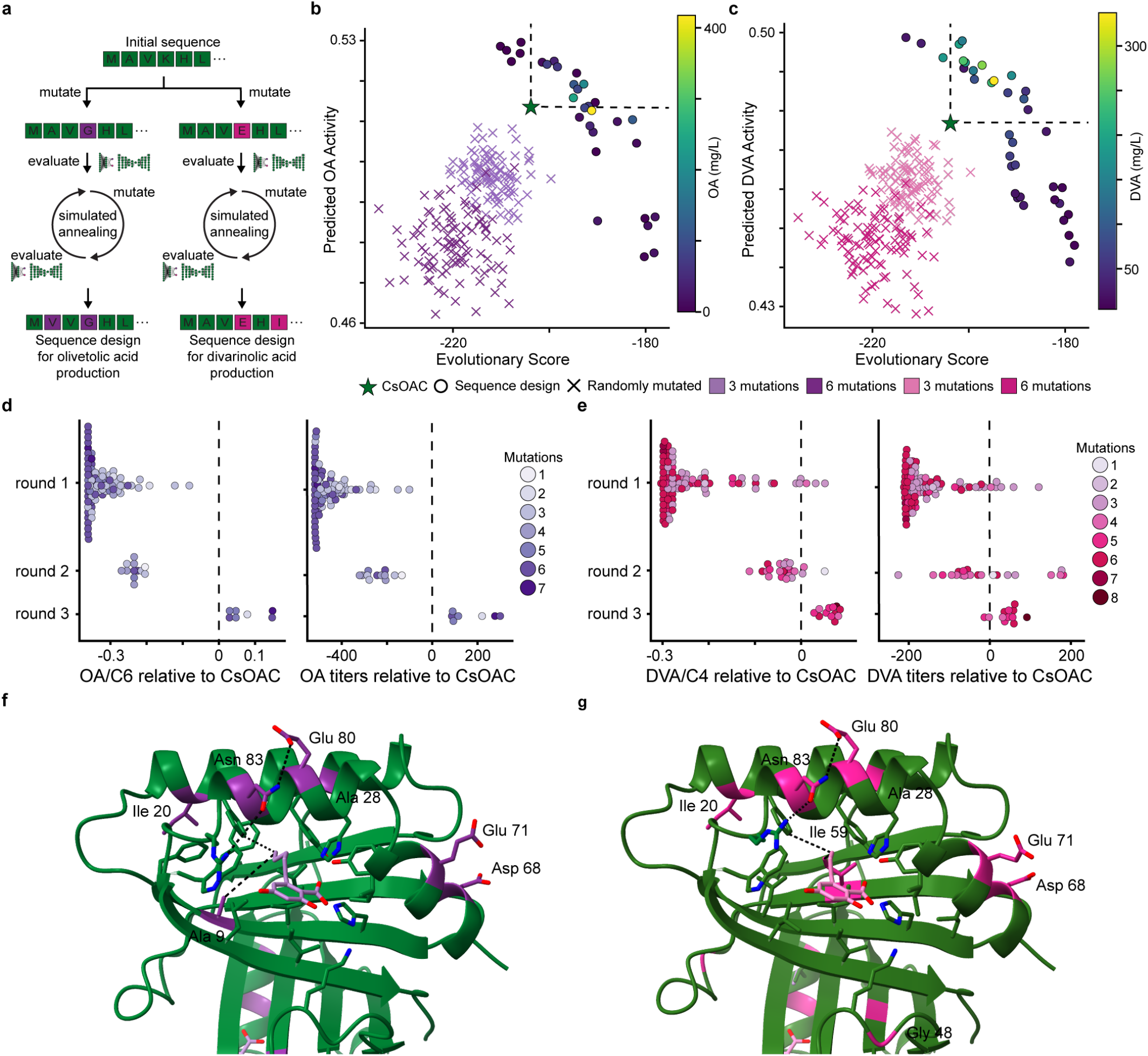
ML-driven evolution of CsOAC activity. **a**, Workflow for generating sequence variant designs via simulated annealing. **b**, Pareto front for sequence designs with 3 or 6 mutations generated via simulated annealing for predicted OA activity from the multi-task neural networks and the evolutionary score from the VAE. We score sequence variants with 3 or 6 randomly sampled mutations to the CsOAC sequence to visualize the separation of our sequence designs from random background. Sequence designs are colored by measured OA titers. **c**, Pareto front for sequence designs with 3 or 6 mutations generated via simulated annealing for predicted DVA activity from the multi-task neural networks and the evolutionary score from the VAE. We score sequence variants with 3 or 6 randomly sampled mutations to the CsOAC sequence to visualize the separation of our sequence designs from random background. Sequence designs are colored by measured DVA titers. **d**, OA selectivity relative to CsOAC and OA titers relative to CsOAC per round of ML-driven evolution. **e**, DVA selectivity relative to CsOAC and DVA titers relative to CsOAC per round of ML-driven evolution. **f**, Best sequence design with 7 mutations relative CsOAC for OA production. Mutations are indicated in purple. **f**, Best sequence design with 8 mutations relative CsOAC for DVA production. Mutations are indicated in pink.

Across three rounds of this iterative design-build-test-learn cycle, both desired product titers and pathway selectivity increased (Figure 3d-e, Supplementary Figure 13). The sequence variant CsOAC1.295, containing six mutations relative to wildtype CsOAC, increased OA titers by 300.28 mg/L and OA selectivity by 0.15 relative to CsOAC (Figure 3d). Similarly, CsOAC1.285, containing eight mutations, increased DVA titers by 91.51 mg/L and DVA selectivity by 0.07 relative to CsOAC (Figure 3e). Notably, VAE evolutionary scores also increased across rounds despite the growing mutational distance from CsOAC, indicating that sequence designs simultaneously improved predicted function and evolutionary plausibility. This result is notable given that randomly sampled multi-mutant sequences rarely achieved evolutionary scores greater than CsOAC, with no sequences containing six or more mutations exceeding CsOAC among 500,000 samples (Supplementary Figure 14). We further refined sequence design objectives and constraints throughout the campaign based on experimental outcomes and model predictions, progressively focusing exploration toward regions of sequence space predicted to support both activity and evolutionary plausibility (Supplementary Figure 11-16, Supplementary Table 8).

The highest-performing OA- and DVA-producing variants accumulated different mutations, revealing distinct mutational signatures associated with improved production of each product (Figure 3f-g). The OA-producing variants were characterized by mutation L9A, whereas the DVA-producing variants were characterized by mutation V59I. We therefore examined these mutations in greater detail to understand their contributions to activity and selectivity. In the OA lineage, experimental characterization showed that L9A combined with T68D recapitulated most of the improvement in OA titers observed in the best-performing variants (Supplementary Figure 17). These results suggest that L9A contributes directly to enhanced OA production, potentially by improving active-site compatibility with the native C6-derived intermediate.

In contrast, the DVA lineage was characterized by mutations at V59. Round 2 characterization showed that V59I increased DVA selectivity without substantially increasing DVA titers (Supplementary Figure 18). Targeted mutagenesis further supported this interpretation: V59L, V59I, and V59M all increased DVA/DVO selectivity, with larger hydrophobic substitutions generally producing stronger effects, while normalized DVA titers remained near baseline (Supplementary Figure 19). Together, these results suggest that V59 primarily controls side-chain selectivity rather than catalytic output. Increased DVA titers emerged only when V59I was combined with additional mutations outside the active site that were shared among high-performing OA- and DVA-producing variants (Figure 3d-g). Several of these mutations cluster in α-helices that form the entrance to the catalytic cavity. D83N and G80E introduce a new salt bridge that may stabilize one helix, whereas D71E and T68D increase the negative surface charge of a neighboring helix and may alter local flexibility. These observations suggest that active-site mutations primarily govern substrate selectivity, while second-shell mutations near the catalytic cavity enhance productive cyclization and overall catalytic performance.

### Engineered CsOACs produce major and minor cannabinoids in vivo

We introduced the OA-optimized variant CsOAC1.295 and the DVA-optimized variant CsOAC1.285 into an engineered *Y. lipolytica* production strain (SB3829) containing the downstream pathway enzymes required to convert OA into THCA and DVA into THCVA, enabling end-to-end biosynthesis of major and minor cannabinoids *in vivo* (Figure 4a-b, Supplementary Table 1-3, Supplemental Figure 2). Strains expressing CsOAC1.295 produced 108 mg/L THCA, alongside 143 mg/L OA and byproducts 55 mg/L OL and 50 mg/L PDAL (Figure 4c). Strains expressing CsOAC1.285 produced 78 mg/L THCVA, together with 289 mg/L DVA and byproducts 54 mg/L DVO and 249 mg/L VDAL (Figure 4d). To our knowledge, this represents the first demonstration of THCA and THCVA production in *Y. lipolytica*. These titers correspond to 13.5-fold (THCA) and 16.25-fold (THCVA) increases relative to previously reported engineered *Saccharomyces cerevisiae* systems^16^ (Supplementary Table 9). Although OA is an intermediate, CsOAC1.295 achieved OA titers 15.58-fold higher than prior reports in *Y. lipolytica* strains expressing only upstream pathway enzymes, indicating substantial improvement in precursor supply even in the presence of downstream pathway flux^18^ (Supplementary Table 9).

**Figure 4:**
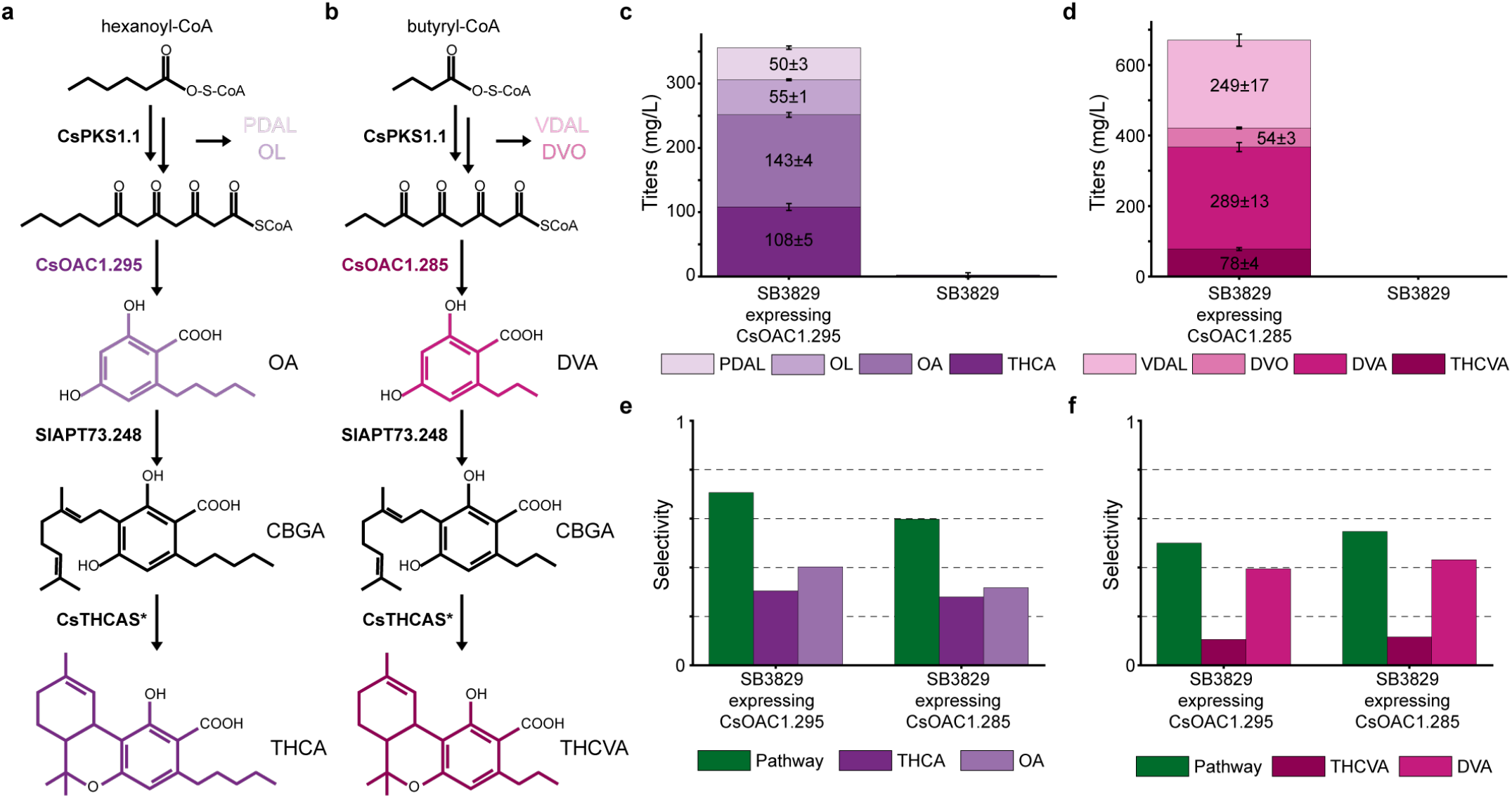
Engineered CsOACs produce major and minor cannabinoids *in vivo*. **a**, THCA production in genetically modified *Y. lipolytica* strain SB3829 (Supplementary Table 1-3). **b**, THCVA production in genetically modified *Y. lipolytica* strain SB3829 (Supplementary Table 1-3). Both pathways include the tetraketide synthase CsPKS1.1 from *C. sativa* that catalyzes the reaction for producing the unstable tetraketide intermediate that can become undesirable byproducts (a) PDAL and OL or (b) VDAL and DVO. Our engineering sequence variants (a) CsOAC1.295 or (b) CsOAC1.285 cyclize the linear tetraketide intermediate to produce (a) OA or (b) DVA. The aromatic prenyltranferase SlAPT73.248 and CsTHCAS* function independently of side chain selectivity to produce (a) THCA or (b) THCVA. **c**, Titers of desired products THCA and OA and byproducts OL and PDAL for our strain SB3829 with and without CsOAC1.295. **d**, Titers of desired products THCVA and DVA and byproducts DVO and VDAL for our strain SB3829 with and without CsOAC1.285. **e**, Pathway selectivity (THCA+OA)/(THCA+OA+OL+PDAL), THCA selectivity THCA/(THCA+OA+OL+PDAL), and OA selectivity OA/(OA+OL+PDAL) for our engineered strains. **f**, Pathway selectivity (THCVA+DVA)/(THCVA+DVA+DVO+VDAL), THCVA selectivity THCVA/(THCVA+ DVA +DVO+VDAL), and DVA selectivity DVA/(DVA+DVO+VDAL) for our engineered strains.

We next quantified pathway selectivity within SB3829. The selectivity for desired products (OA + THCA) relative to byproducts was 0.71 for CsOAC1.295, whereas selectivity for (DVA + THCVA) was 0.548 for CsOAC1.285 (Figure 4e-f). Consistent with design objectives, OA selectivity was 1.27-fold higher in CsOAC1.295 compared to CsOAC1.285, while DVA selectivity was 1.09-fold higher in CsOAC1.285 (Figure 4e). DVA selectivity for SB3829 expressing CsOAC1.285 (engineered for DVA production) was 1.09-fold higher than SB3829 expressing CsOAC1.295 (Figure 4f). Despite these substantial improvements, substantial pools of OA and DVA remained unconverted to THCA and THCVA indicating a downstream bottleneck requiring future engineering.

## Discussion

Cannabinoids comprise a structurally diverse class of bioactive molecules with important therapeutic potential, and expanding access to both major and minor cannabinoid chemotypes remains a central challenge for biomanufacturing and drug discovery. Here, we show that machine learning-guided enzyme engineering can relieve a key bottleneck in cannabinoid biosynthesis by improving the activity and substrate selectivity of olivetolic acid cyclase (OAC). Across three rounds of ML-driven evolution, we identified CsOAC variants that increase production of both OA and DVA, enabling downstream biosynthesis of the major cannabinoid precursor THCA and the minor cannabinoid precursor THCVA in *Yarrowia lipolytica* at titers exceeding prior microbial systems. These results position CsOAC as a critical control point for pathway performance and demonstrate that targeted improvement of an early enzymatic step can propagate to increased pathway flux and altered product distributions. More broadly, the ability to reprogram CsOAC substrate preference suggests a path toward expanding cannabinoid chemical diversity through the production of both naturally occurring minor cannabinoids and, potentially, non-natural analogues derived from alternative precursors.

Engineering cannabinoid cyclases presents two related challenges. First, cannabinoid-producing activity appears to be rare in nature. Despite screening 40 homologous polyketide cyclases, none produced detectable levels of OA or DVA. This observation is consistent with the limited distribution of cannabinoid biosynthesis in nature and suggests that homolog screening alone may be insufficient for identifying enzymes with the desired activity or selectivity. Second, the CsOAC sequence-function landscape appears highly constrained and epistatic. Consistent with prior studies and our initial mutagenesis libraries, single substitutions rarely yielded measurable gains in activity or selectivity, whereas the highest-performing variants identified here contained multiple coordinated substitutions. Together, these observations indicate that productive CsOAC variants are unlikely to be found either through exploration of natural sequence diversity or through iterative rounds of laboratory evolution, highlighting the need for approaches capable of identifying beneficial combinations of interacting mutations.

To address this challenge, we combined a supervised multi-task predictor with a generative evolutionary prior. The supervised model learned pathway-relevant sequence-function relationships directly from measured product and byproduct distributions, enabling optimization toward experimentally observed pathway outcomes. In principle, the model can assign functional scores to any sequence, including variants far outside the mutational regime represented in the training data. However, confidence in these predictions decreases as sequence designs move further from experimentally characterized regions of sequence space. The evolutionary VAE provides a complementary constraint by capturing evolutionary patterns across the broader polyketide cyclase family and biasing sequence proposals toward variants consistent with natural sequence diversity. Rather than predicting function directly, the VAE restricts exploration to regions of sequence space more likely to retain structural integrity and catalytic competence. Together, these models balance functional innovation and evolutionary conservation, enabling efficient exploration of protein sequence space and providing a general framework for engineering complex protein functions from limited experimental data.

Our results further emphasize the importance of multi-objective optimization for enzymes embedded in metabolic pathways. For CsOAC, improved performance requires not only increased catalytic turnover but also suppression of competing reactions that divert flux through derailment products. Because the tetraketide product is chemically unstable, productive cyclization must kinetically outcompete decarboxylative-aldol condensation, which forms OL/DVO. By jointly modeling desired products and byproducts, our approach captures these competing fluxes explicitly and selects for variants that improve net pathway efficiency rather than redistributing carbon among undesired outcomes. This formulation is broadly applicable to pathway engineering problems in which intermediate stability and flux partitioning constrain overall yield.

Analysis of top-performing variants provides initial insight into the structural determinants of CsOAC function. Mutations such as L9A and V59I, located within the substrate-binding pocket, are consistent with direct effects on substrate accommodation and side-chain selectivity, particularly in distinguishing C6- and C4-derived intermediates. In contrast, several enriched mutations occur in helices at the entrance to the catalytic cavity, where they may alter local packing or electrostatics and thereby influence conformational dynamics associated with intermediate binding or release. Together, these observations suggest that CsOAC activity and selectivity are governed by a combination of active-site geometry and second-shell or entrance-region effects. While these hypotheses are consistent with the observed mutational patterns, resolving their mechanistic basis will require further targeted structural and kinetic characterization.

Despite substantial improvements in precursor formation and pathway selectivity, downstream conversion of OA and DVA to THCA and THCVA remains incomplete, indicating that additional pathway steps now limit overall production. This shift in bottleneck is consistent with a hierarchical view of pathway optimization, in which relieving one constraint reveals the next limiting process. Further improvements will likely require coordinated engineering of prenyltransferases, oxidocyclases, precursor supply, and host metabolic state. In this context, the present work demonstrates how targeted optimization of a single enzymatic node can be used to iteratively expose and address subsequent limitations in a complex biosynthetic network.

More broadly, this study illustrates a strategy for engineering enzymes whose function cannot be readily decoupled from pathway context. For systems involving unstable intermediates or competing side reactions, *in vitro* assays may not faithfully capture determinants of *in vivo* performance. By training directly on pathway-level measurements, our framework evaluates variants under conditions that reflect their functional role within the full biosynthetic system. This framework is particularly valuable for metabolic engineering applications in which enzyme activity, flux distribution, and cellular context are tightly coupled. In parallel, tuning CsOAC substrate selectivity provides a route to expanding accessible cannabinoid chemical space. Although this work focuses on C6- and C4-derived pathways, the same approach could be extended to enable acceptance of alternative fatty acid–derived precursors, supporting biosynthesis of cannabinoid analogues with varied side-chain structures. Because side-chain identity strongly influences physicochemical and pharmacological properties, this capability may facilitate systematic exploration of structure–activity relationships across both abundant and less characterized cannabinoid classes.

Advances in synthetic biology are making it increasingly routine to transplant complex biosynthetic pathways from plants and other organisms into engineered microbial hosts. The challenge is no longer simply reconstructing these pathways, but learning how to control them. By coupling pathway-level measurements with machine learning-guided enzyme engineering, it becomes possible to learn how individual enzymes shape the production of complex molecules and to rationally redirect pathway outputs toward desired products. As larger datasets accumulate across diverse biosynthetic systems, such approaches may transform metabolic engineering from a process of manual optimization into one of data-driven pathway design, expanding access to valuable natural products, minor metabolites, and entirely new chemical matter.

## Methods

### Initial library creation

We accumulated an initial sequence variant library of 151 olivetolic acid cyclase (OAC) variants plus the wild-type (WT) enzyme from *C. sativa* using three complementary design strategies. First, we screened a panel of 40 CsOAC homologs. None of the homologs produced OA or DVA above the analytical limit of detection. Second, we generated a single site-saturation mutagenesis library to systematically sample amino-acid substitutions at targeted positions. Third, we created recombination variants by swapping secondary-structure elements from selected homologs into the CsOAC scaffold. These included a *C. sativa* homolog that produced low levels of OA/DVA *in vitro* but not *in vivo*, and a *Rhododendron dauricum* homolog that natively produces the C2 (1-carbon tail) analogue of OA (Supplementary Figure 3).

### In vivo characterization

Plasmids were transformed into strain SB-3829 by electroporation. Colonies were patched onto YPD + 1 g/L hygromycin plates [YPD = 10 g/L Bacto Yeast Extract (Gibco, Cat. No. 212750), 20g/L Bacto Peptone (Gibco, Cat. No. 211677), 22 g/L Dextrose Monohydrate (Bakers Authority, Cat. No. D-7950), and 20 g/L Difco Agar (BD Biosciences, Cat. No. 214010)]. These patches were used to inoculate seed cultures in 500 uL YDKMU13 + 0.1 g/L hygromycin [YDKMU13 = 1.71 g/L YNB (Sunrise Science, Cat. No. 1500-5KG), 66 g/L (6%) Dextrose Monohydrate (Bakers Authority, Cat. No. D-7950), 13 g/L Hy-Express System I (K615) (Kerry, SAP Code – 20070615), 100 mM (19.5 g/L) MES Hydrate (Sigma-Aldrich, Cat. No. M2933), and 11.5 g/L Urea (Fisher Chemical, U15-500); adjust pH to 6.5 using 8N KOH (Thermo Scientific, Cat. No. 380620020)] in 96 deep well (pyramid bottom) plates and grown overnight in a high-speed shaker (990 rpm) at 30°C. Fermentations were started by inoculating 500 uL YDKMU13 + 0.1 g/L hygromycin with 5 uL of the seed culture. At 24h and 30h of fermentation, cultures were spiked with 25 uL 100 mM butyric acid or 25 uL 100 mM hexanoic acid. At 48h of fermentation, the entire culture was quenched with 500 uL Quench solution (0.2 g/L internal standard (3,5-diisopropyl-2-hydroxybenzoic acid 98%) in reagent alcohol) and analyzed for cannabinoids and relevant intermediate metabolites.

### Data curation for VAE

We curated 14,722 natural sequences related to olicetolic acid cyclase (OAC) from the protein database UniRef90^43^ with the Hidden-Markov model homology search tool Jackhmmer^44^ to obtain a multiple-sequence alignment (MSA) containing proteins related to CsOAC with the default number of iterations (N=5). We removed sequences less than 75% of the length of CsOAC^45^, removed sequences with an amino acid repeating 10 times in a row, and removed positions of the MSA not corresponding to CsOAC^46^. We reweighted the remaining sequences with neighbors classified as having a Hamming distance/length of sequence greater than 0.8 to reduce phylogenetic bias from uneven sampling^45,47,48^. We randomly split the MSA sequences into a 90% training set and a 10% validation set.

### Variational autoencoder training and inference

The VAE is trained by maximizing a modified version of the Evidence Lower Bound (ELBO) that effectively minimizes the Kullback–Leibler divergence between the variational approximation and the true posterior distribution^45^ as shown with equation (1):

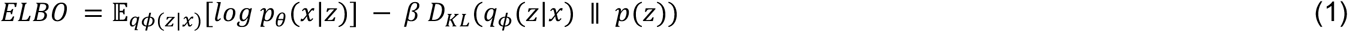

The first term can be considered a reconstruction loss that is computed using cross-entropy between an input one hot-encoded sequence and output likelihoods. β is the weight for the Kullback–Leibler divergence term *D*_*KL*_ with a prior distribution *p*(*z*) of *N*(0, *I*). Model architecture and training hyperparameters are included in Supplementary Table 6.

To compute a VAE-based evolutionary score for each sequence *x* = (*x*_1_, …, *x*_*L*_), we used the VAE decoder to compute the summed reconstruction cross-entropy across sequence positions. Because the VAE uses stochastic latent sampling, we obtained deterministic scores by decoding from the mean of the approximate posterior, *z*_*μ*_, rather than from a sampled latent variable. The reconstruction loss was obtained with equation (2):

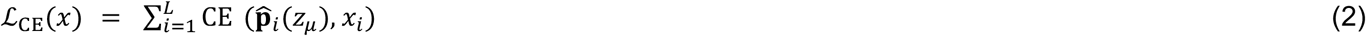

where 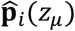 is the decoder-predicted categorical distribution at position *i* conditioned on *z*_*μ*_. We defined the VAE evolutionary score as the negative reconstruction loss so that sequences with higher VAE-assigned likelihoods receive higher evolutionary scores as shown with equation (3):

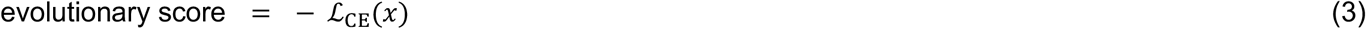

### Training ensemble of multi-task multi-layer perceptron models

We trained an ensemble of 100 multi-task multi-layer perceptron (MLP) regression models to predict metabolite production from olivetolic acid cyclase (OAC) variant sequences. Each model jointly predicted normalized assay readouts corresponding to each experimental condition (combinations of strain, feedstock, measurement timepoints, and media), with four outputs per condition capturing product/byproduct ratios and yields (e.g., OA/C6, OA/OL, OA/(OA+OL), and (OA+OL)/PDAL for hexanoic-acid feeding and analogous DVA metrics for butanoic-acid feeding). Because not all variants were measured for all conditions, missing label values were encoded and excluded from the loss using an observation mask. Labels were max-normalized per output dimension to avoid upweighting outputs with larger numerical ranges. During training, we minimized a masked, label-weighted mean-squared error (MSE) objective averaged over observed labels in each minibatch using equation (4):

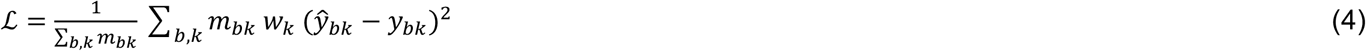

where *m*_*bk*_ ∈ {0,1} indicates whether output *k* is observed for sequence *b*, and *w*_*k*_ are fixed per-output weights used to down-weight ratio terms relative to primary readouts (Supplementary Table 5).

Each round, we clustered experimentally characterized sequences by one-hot encoding each position and applying K-means to the resulting flattened vectors. Then, we used the resulting cluster labels to stratify train/validation/test splits so that each split contained representatives from across sequence space so that validation loss during training reflected performance across the diverse sequence space. This split was fixed across all ensemble members, and each model was trained independently and selected based on minimum validation loss. All model architecture choices and training hyperparameters (including any round-specific changes in output dimensionality, hidden widths, loss-weight patterns, and training schedule) are summarized in Supplementary Table 5.

### Computing functional score with multi-task multi-layer perceptron models

We computed a single functional score to rank candidate sequence variants of CsOAC from the ensemble’s multi-output predictions. For each sequence, we evaluated all 100 MLPs to obtain predicted values for every output dimension, then collapsed these predictions to a scalar by focusing on target outputs relevant to the optimization objective (olivetolic acid or divarinic acid production) and averaging them. In round 1, we used the ensemble mean prediction for each output and defined the score as the mean of the two selected outputs with equation (5):

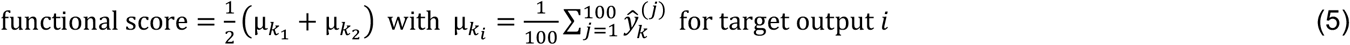

Here, 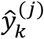 is the prediction from ensemble member *j* for output *k* and (*k*_1_, *k*_2_) are the target outputs used for scoring (Supplementary Table 7).

In subsequent rounds, we used a more conservative lower confidence bound (LCB) across ensemble outputs given the success of this in literature^49^, computed as the 5th percentile of model predictions for each output, and averaged the same target outputs with in equation (6):

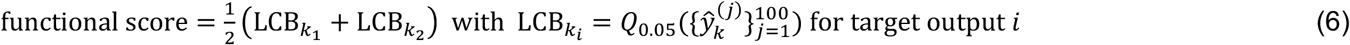

Here, 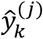 is the prediction from ensemble member *j* for output *k, Q*_0.05_denotes the 5th percentile, and (*k*_1_, *k*_2_) are the two target outputs used for scoring (Supplementary Table 7).

### Generating designs via simulated annealing

We generated CsOAC sequence variant designs via simulated annealing by adapting code from prior work^49^. We designed sequence variants with a fixed number of substitutions to maximize a composite reward with two components: (i) a VAE evolutionary score and (ii) a functional score from a multi-task MLP ensemble. At each step, we proposed a new sequence by perturbing the current mutant. The number of mutation edits per proposal was sampled from *n* ∼ Poission(*λ* = mut_rate) and constrained so that at least one edit was made while the total number of substitutions remained fixed (Supplementary Table 7-8). Proposals were generated by removing *n* existing substitutions and introducing *n* new substitutions at allowable positions, maintaining exactly *N*_mut_ substitutions per design. Candidate mutations were applied as point substitutions at allowable positions (N- and C-termini residues were fixed for cloning and amino-acid size constraints were applied at active-site positions during round 3; Supplementary Table 8, Supplementary Figure 16).

Because the VAE and MLP-derived scores had different numeric scales, we min–max normalized each component prior to multi-objective combination. For each design setting, normalization bounds were estimated empirically by running single-objective simulated annealing using either the VAE evolutionary score alone or the MLP ensemble functional score. Normalized scores were computed with equation (7) and equation (8):

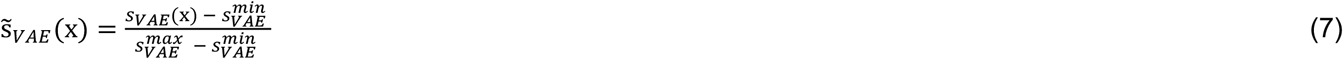

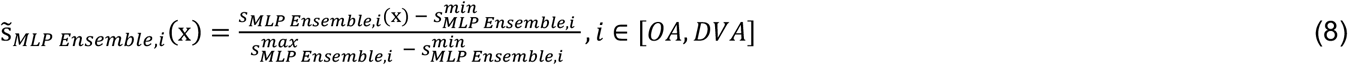

For Pareto design, we combined normalized scores with equation (9):

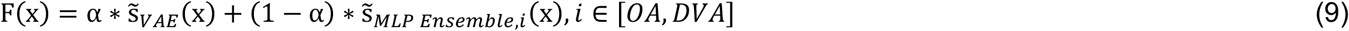

where *α* ∈ [0,1] was swept over a grid to generate variants spanning trade-offs between evolutionary plausibility and predicted function. For rounds 2 and 3, we also performed utopia design by minimizing the Euclidean distance to the utopia point (1,1) in normalized score with equation (10):

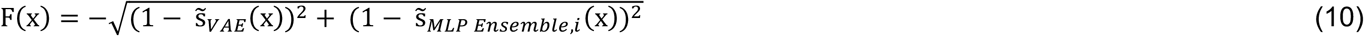

Mutations that enhanced function were automatically accepted while all other mutations were accepted with a probability calculated with equation (11):

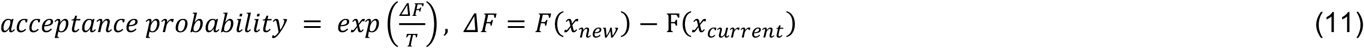

where T decreases with a logarithmic temperature gradient selected to have an initial acceptance probability near 40% for mutations that do not increase function to encourage early exploration and approaches 0% towards the end of the simulation. Additional parameters are indicated in Supplementary Table 8. Each round, we clustered generated designs using K-means++ to select diverse sequence designs for experimental characterization.

### OAC evolution details

Sequence design objectives and constraints were refined across rounds based on experimental outcomes and model predictions. Following round 1, we observed that variants with both functional and evolutionary scores exceeding those of CsOAC were more likely to retain enzyme activity. Consequently, rounds 2 and 3 incorporated both Pareto-front sampling and utopia design strategies to enrich for variants with simultaneously high functional and evolutionary scores (Figure 3b-c, Supplementary Figure 11-12). For rounds 2 and 3, terminal residue constraints were relaxed to permit mutations at N-terminal positions predicted to be beneficial by the design models (Supplementary Figure 15). For round 3, an additional structural constraint was introduced for OA design, restricting substitutions that reduced active-site pocket volume. Active-site residues were defined as residues within 4 Å of docked OA, and candidate designs violating this constraint were excluded (Supplementary Figure 16, Supplementary Table 8).

## Data Availability Statement

The work described uses multiple rounds of characterization of sequence variants for CsOAC variants that can be found on Github (https://github.com/RomeroLab/ml-guided-cannabinoid-biosynthesis/tree/main/data).

## Code Availability Statement

All code is available on the public, open-source GitHub repository: https://github.com/RomeroLab/ml-guided-cannabinoid-biosynthesis under the Apache 2.0 license.

