## Supplementary Information for "Machine learning-guided olivetolic acid cyclase engineering enables tailored cannabinoid biosynthesis in yeast"

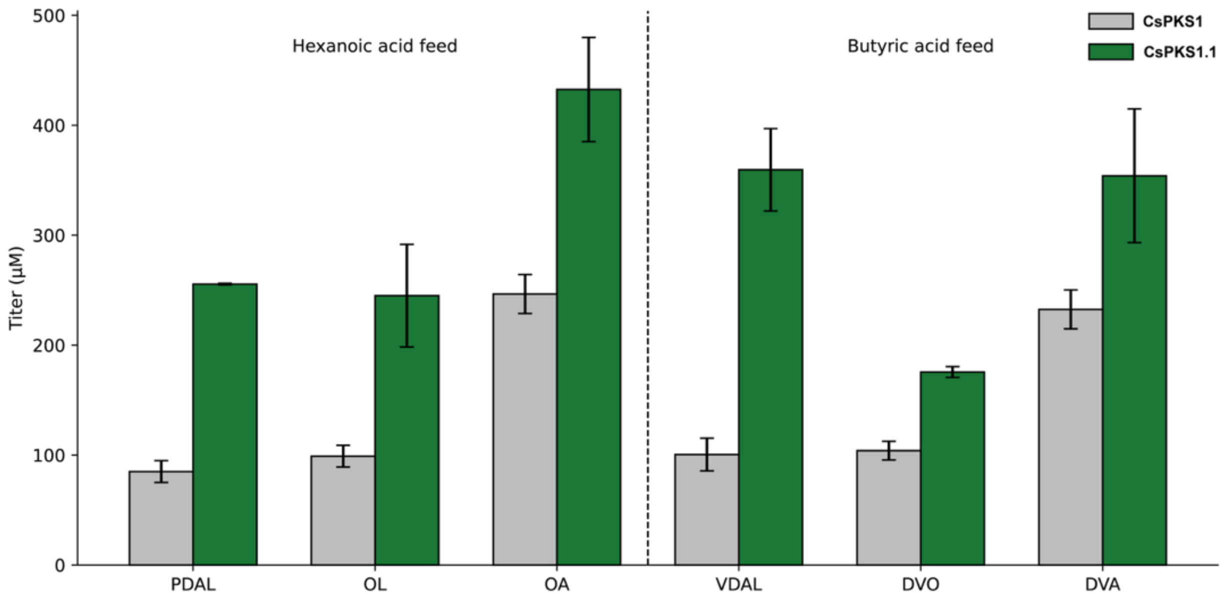


Supplementary Figure 1: CsPKS1.1 increases polyketide-derived product titers in engineered *Yarrowia lipolytica*. Average titers of hexanoic acid-derived products, pentyl diacetic acid lactone (PDAL), olivetol (OL), and olivetolic acid (OA), and butyric acid-derived products, varinic diacetic acid lactone (VDAL), divarinol (DVO), and divarinic acid (DVA), produced by *Y. lipolytica* strains expressing genomically integrated AtHCS2 and CsOAC and plasmid-encoded CsPKS1 or CsPKS1.1. Colonies were grown for 48 hr in YPD containing 6% glucose, 5 mM $MgSO_4$, and 1 mg/mL hygromycin, then used to inoculate assay cultures supplemented with 2.5 mM hexanoic acid or butyric acid. After 24 hr, cultures were supplemented with an additional 2% glucose and 5 mM corresponding fatty acid feed. After 48 h total growth, cultures were quenched and analyzed by LC. Bars represent mean titers from two biological replicates with error bars indicating standard deviation.

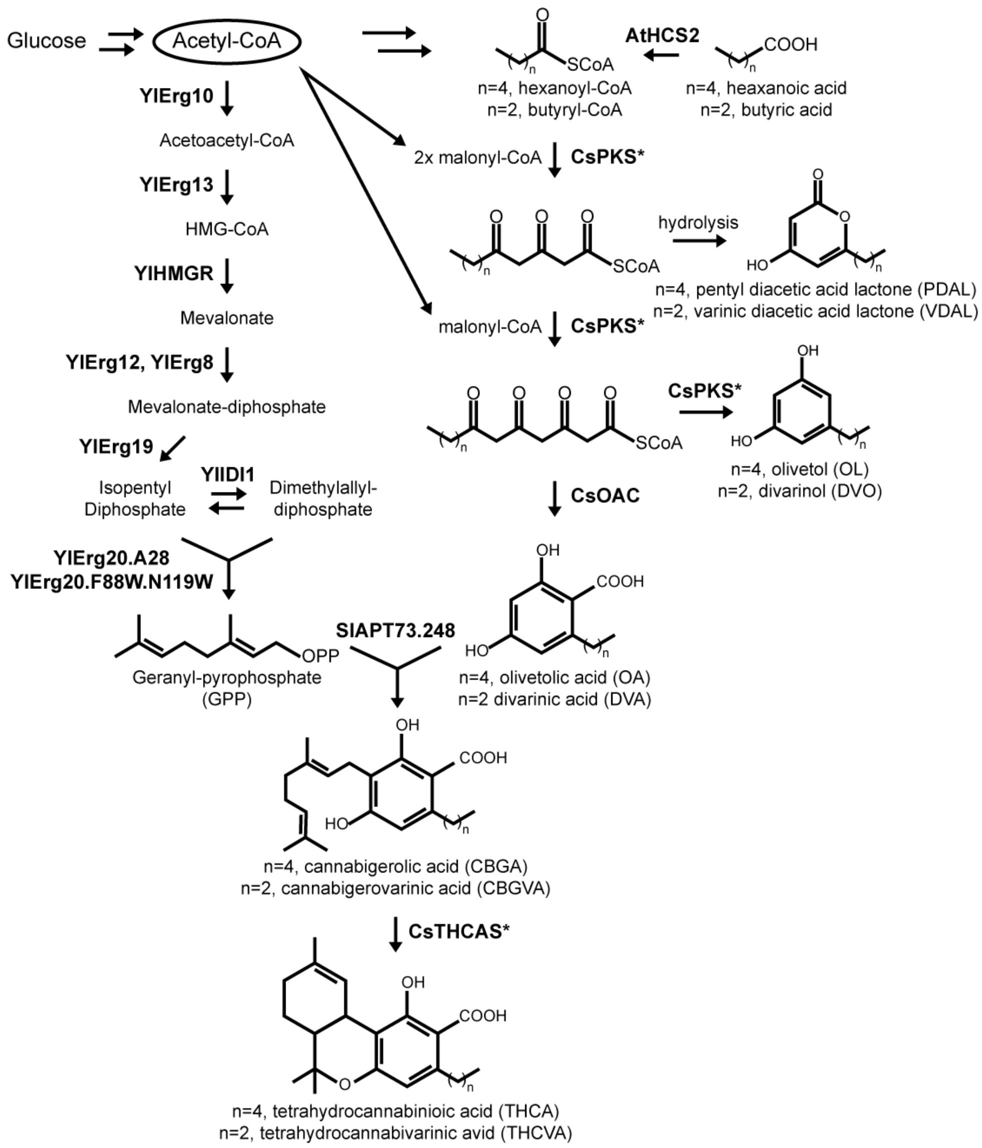


Supplementary Figure 2: Cannabinoid biosynthesis in *Y. lipolytica*. Expression of AtHCS2 enables production of hexanoyl-CoA or butyryl-CoA after from hexanoic acid or butyric acid feeds. Expression of CsPKS* and CsOAC enables production of olivetolic acid (OA) or divarnic acid (DVA) from hexanoyl-CoA or butyryl-CoA. The reactive linear tetraketide intermediate after condensations with two malonyl-CoA units can undergo abiotic hydrolysis to produce undesirable byproduct pentyl diacetic acid lactone (PDAL). The reactive linear tetraketide intermediate after condensations with three malonyl-CoA can be cyclized by CsPKS* to produce undesirable byproduct olivetol (OL). Expression of SlAPT73.248 enables production of cannabigerolic acid (CBGA) and cannabigerovarinic acid (CBGVA) via the prenylation of OA or DVA. The expression of CsTHCAS* enables production of tetrahydrocannabinolic acid (THCA) or tetrahydrocannabivarinic acid (THCVA). Geranyl-pyrophosphate (GPP) was produced with the expression of the genes YlErg10, YlErg13, YlHMGR, YlErg12, YlErg8, YlErg19, YlErg20.F88W.N119W[1] in *Y. lipolytica*[2,3]. Engineered variants of CsPKS and CsTHCAS are indicated with an asterisk (Supplementary Table 1-2).

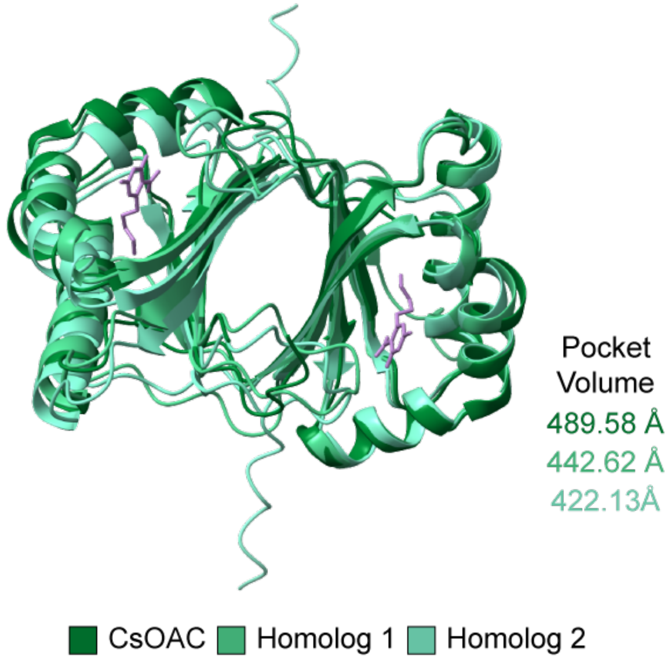


Supplementary Figure 3: Structures of Olivetolic Acid Cyclase and Secondary-Structure Donor Homologs. We obtained the crystal structure for CsOAC bound to olivetolic acid from *Cannabis sativa* (PDB: 5B09). We used Alphafold3 to predict the structures for Homolog 1 from *Cannabis sativa* that produced low levels of OA/DVA *in vitro* (but not in *Yarrowia lipolytica*) and Homolog 2 from *Rhododendron dauricum* that natively produces the 1-carbon tail analogue of olivetolic acid, orsellinic acid. We aligned the predicted homolog structures to the CsOAC crystal structure bound with olivetolic acid (PDB: 5B09) using least-squares fitting of specified atoms between structures using ChimeraX. Pocket volumes were estimated using CAVER software.

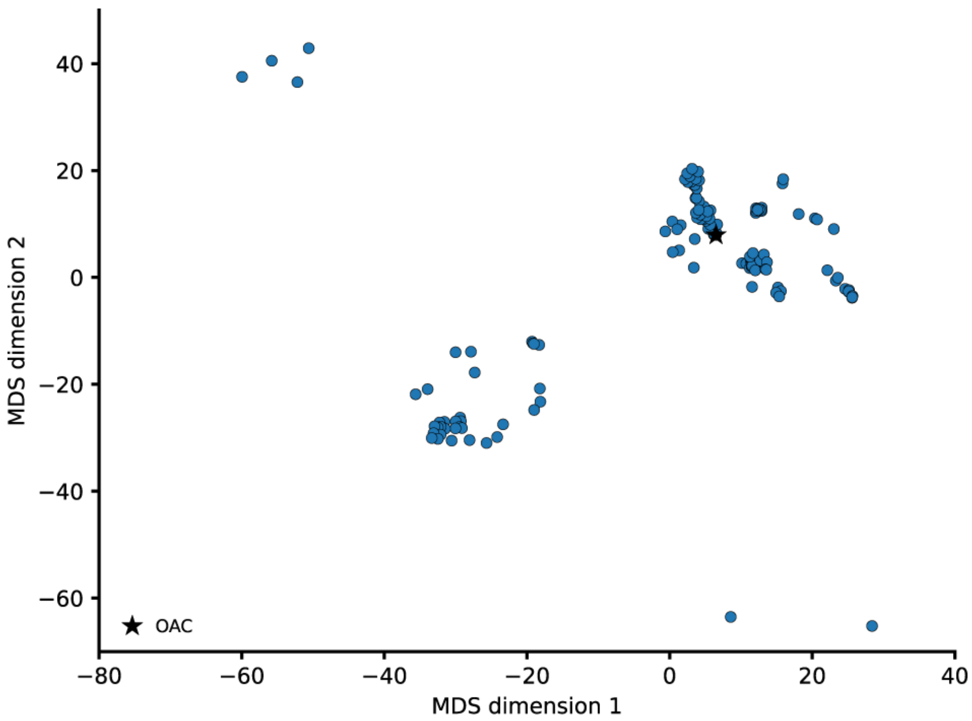


Supplementary Figure 4: Multi-dimensional scaling (MDS) visualization of the sequence from the initial libraries. Distances were computed using pairwise Hamming distance between aligned sequences, counting amino-acid mismatches while ignoring gap positions. The resulting pairwise distance matrix was used as a precomputed dissimilarity input to MDS, such that sequences positioned closer in the 2D embedding space are closer in sequence. Axes correspond to the first two MDS dimensions. Each point represents a sequence variant. The CsOAC sequence is highlighted with a star.

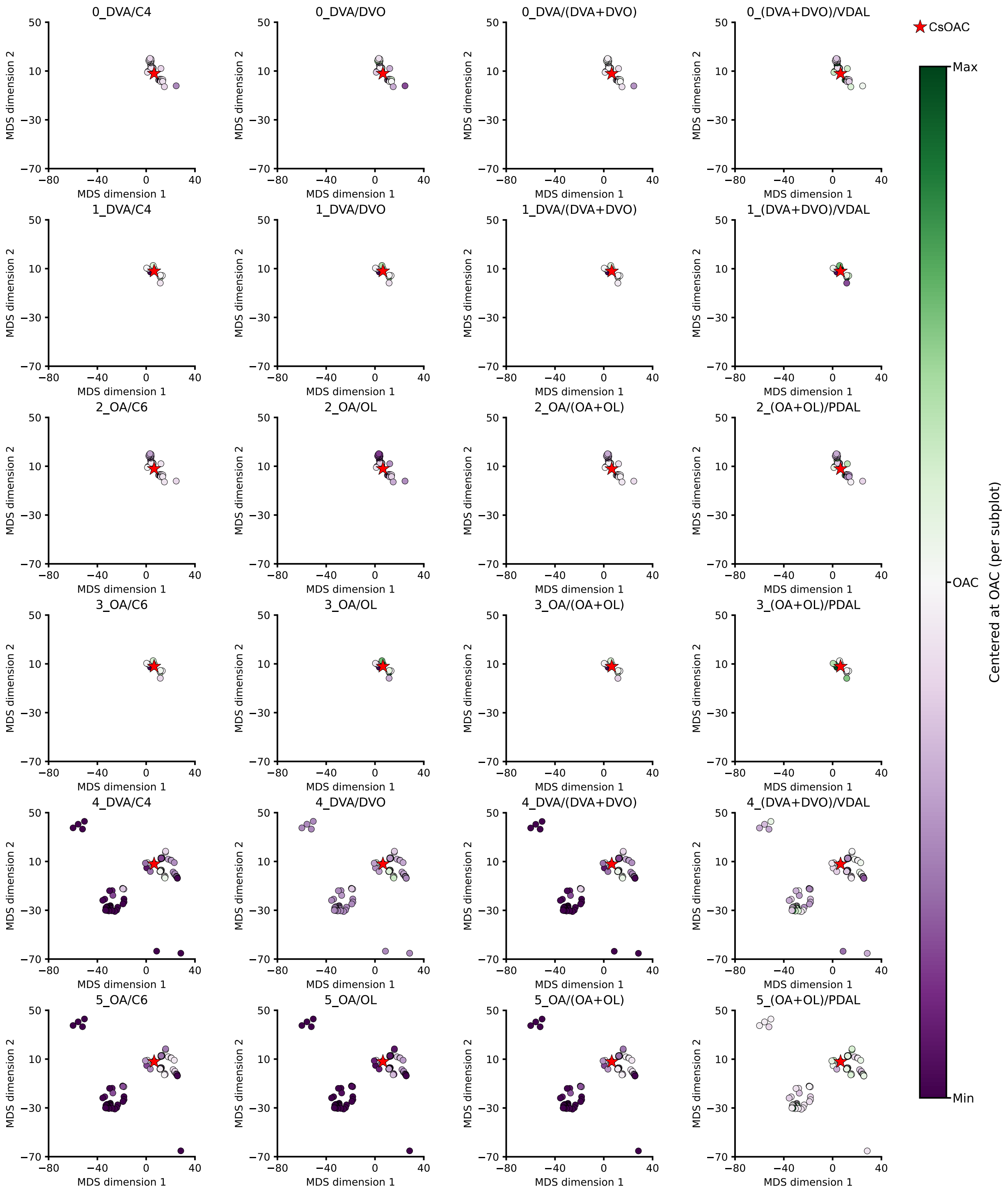


Supplementary Figure 5: Multi-dimensional scaling (MDS) visualization of sequence–function relationships across measured metabolite ratios. Each point represents a sequence variant, colored by the corresponding functional metabolite ratio value centered relative to the CsOAC sequence. The CsOAC sequence is highlighted with a star. Sequence variants are not shown if they were not measured for the corresponding functional metabolite ratio (Supplementary Table 1).

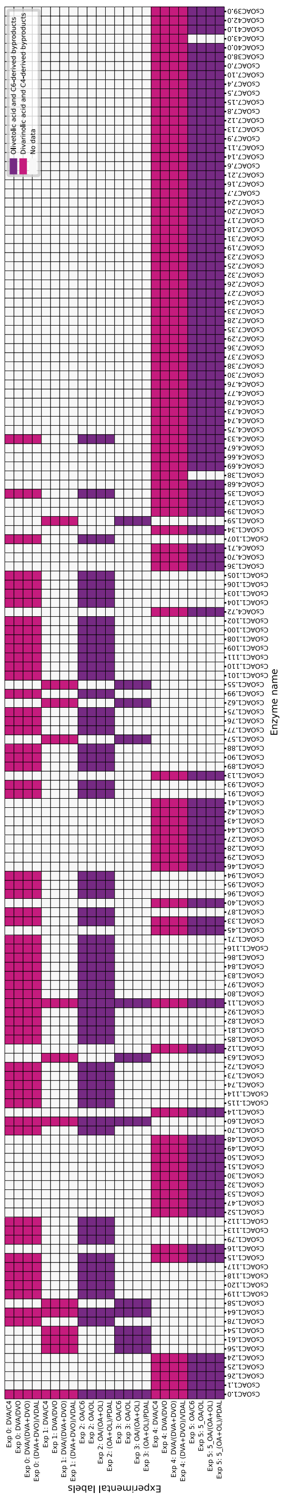


Supplementary Figure 6: Experimental coverage across initial sequence variant library. Heatmap displays experimental measurements obtained for each enzyme variant. Colors indicate assays measuring olivetolic acid (OA) and C6-derived byproducts (purple), divarnic acid (DVA) and C4-derived byproducts (magenta), or missing measurements (gray) illustrating incomplete but some complementary experimental coverage that motivates the multi-task modeling approach.

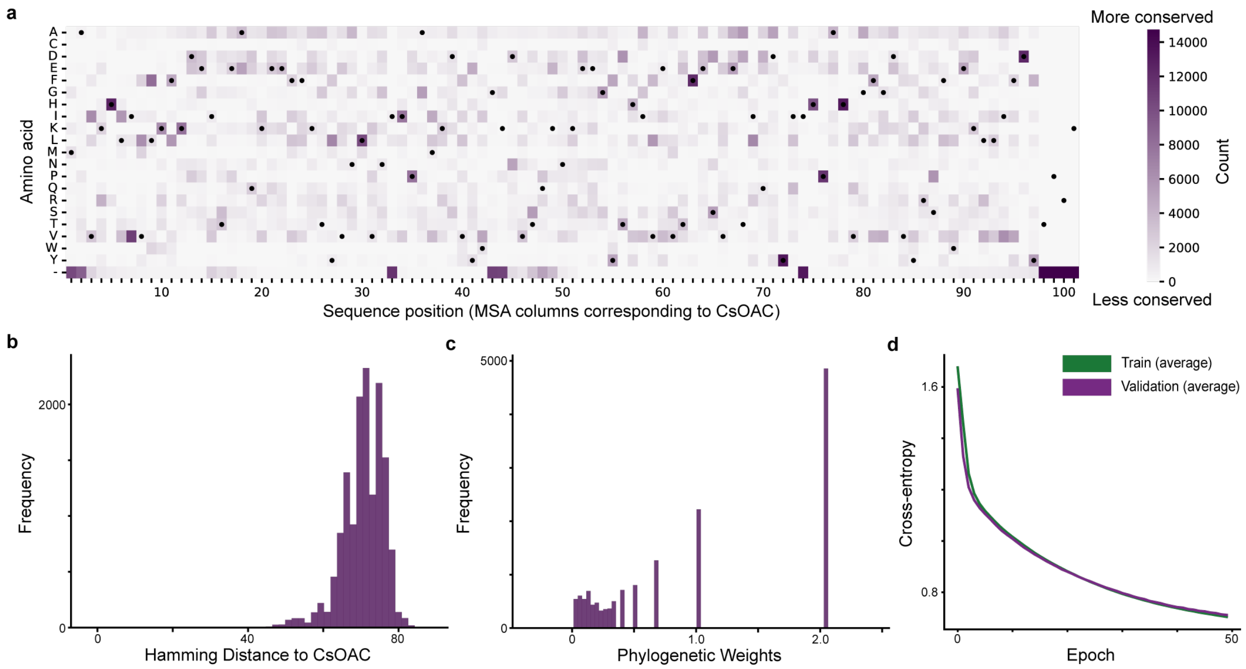


Supplementary Figure 7: VAE pre-training. a, Distribution of sequence identity in curated MSA of natural sequences related to CsOAC used to train VAE. b, Distribution of sequence identity in curated MSA of natural sequences related to CsOAC used to train VAE. c, Distribution of weights for natural sequences in the curated MSA that account for phylogenetic bias in the MSA. d, Training curves for VAE pre-training showing cross-entropy loss.

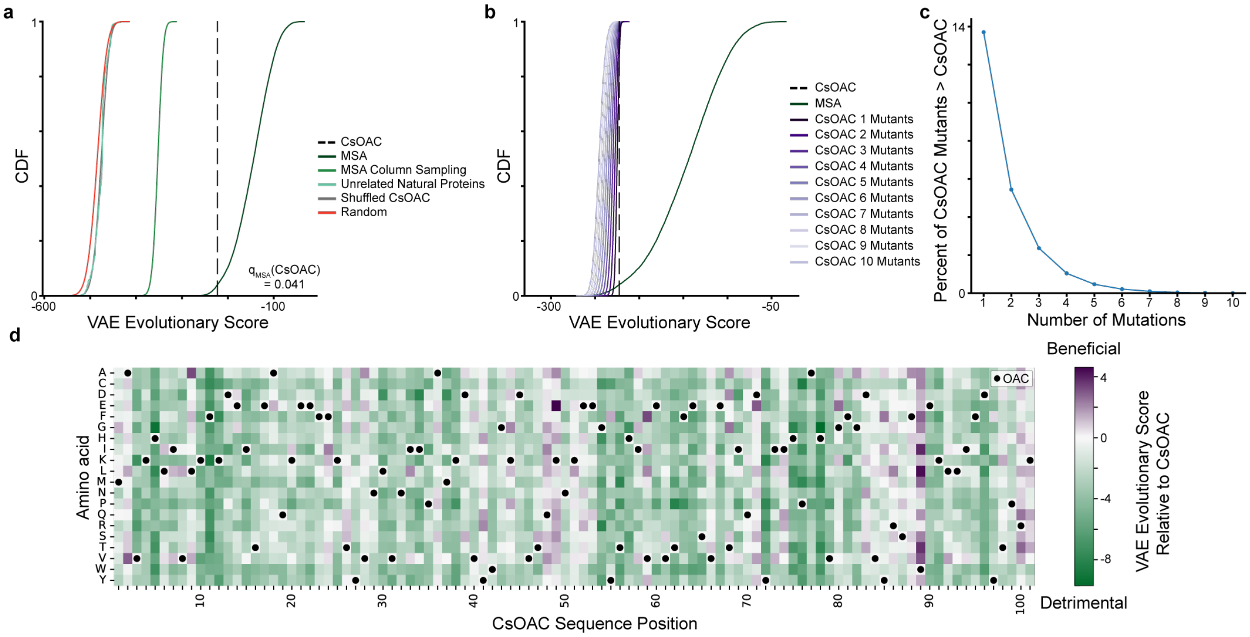


Supplementary Figure 8: Variational autoencoder evolutionary scores characterize statistical constraints of the CsOAC protein family a, Distribution of VAE evolutionary scores across sequence classes used to assess model specificity. Natural CsOAC-related sequences from the multiple sequence alignment (MSA) receive substantially higher scores than control sequence ensembles, including (i) shuffled CsOAC sequences generated by randomly permuting amino acids across non-gap positions while preserving amino-acid composition and gap structure, (ii) column-resampled MSA sequences constructed by independently sampling residues from each alignment column to preserve site-wise frequencies while disrupting higher-order correlations, (iii) unrelated natural proteins matched in sequence length, and (iv) fully random amino-acid sequences. These comparisons demonstrate that the VAE captures higher-order statistical constraints characteristic of the CsOAC family rather than simple amino-acid composition or positional frequencies. b, Evolutionary score distributions for natural CsOAC-related sequences and CsOAC variants containing increasing numbers of mutations relative to wild type. Evolutionary scores progressively decrease as mutational distance from CsOAC increases. c, Fraction of CsOAC mutant sequences with evolutionary scores exceeding wild-type CsOAC. All single and double mutants were exhaustively enumerated and scored, whereas higher mutational regimes were estimated by scoring 500,000 randomly sampled variants per mutation count. The probability of improving evolutionary score declines rapidly with increasing mutational load. d, Position-wise landscape of CsOAC single–amino-acid substitutions colored by the change in VAE evolutionary score relative to wild type, with black markers indicating the wild-type CsOAC sequence.

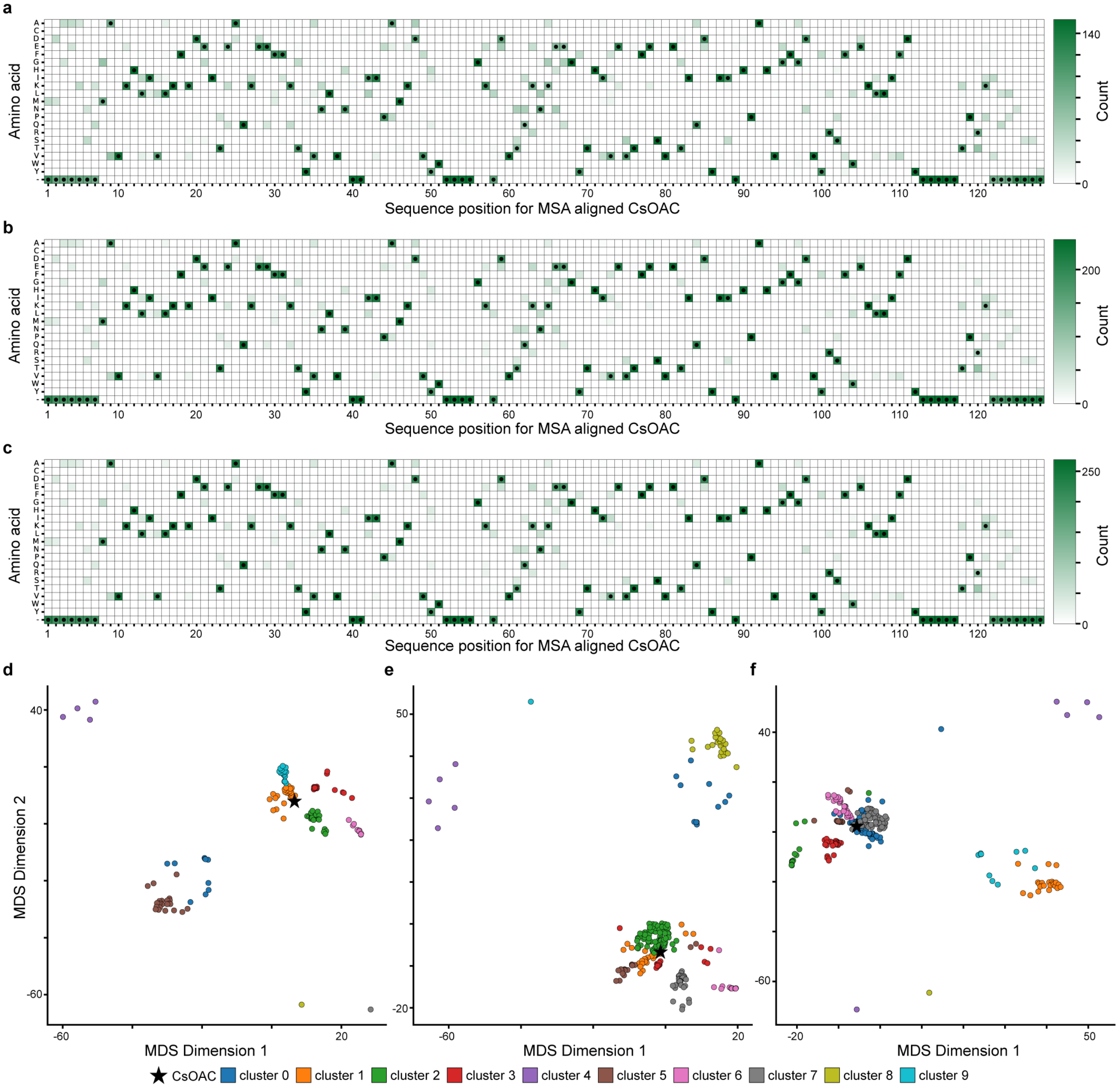


Supplementary Figure 9: Sequence diversity of characterized sequences for training supervised multi-task neural networks. Distribution of sequence identity for sequences used to train multi-task neural networks for round 1 (a), round 2 (b), and round 3 (c). Multi-dimensional scaling (MDS) visualization of the sequence for sequences used to train multi-task neural networks for round 1 (d), round 2 (e), and round 3 (f). Distances were computed using pairwise Hamming distance between aligned sequences, counting amino-acid mismatches while ignoring gap positions. The resulting pairwise distance matrix was used as a precomputed dissimilarity input to MDS, such that sequences positioned closer in the 2D embedding space are closer in sequence. Axes correspond to the first two MDS dimensions. Each point represents a sequence variant. The CsOAC sequence is highlighted with a star. Sequences are colored by clusters used to construct training, validation, and test sets.

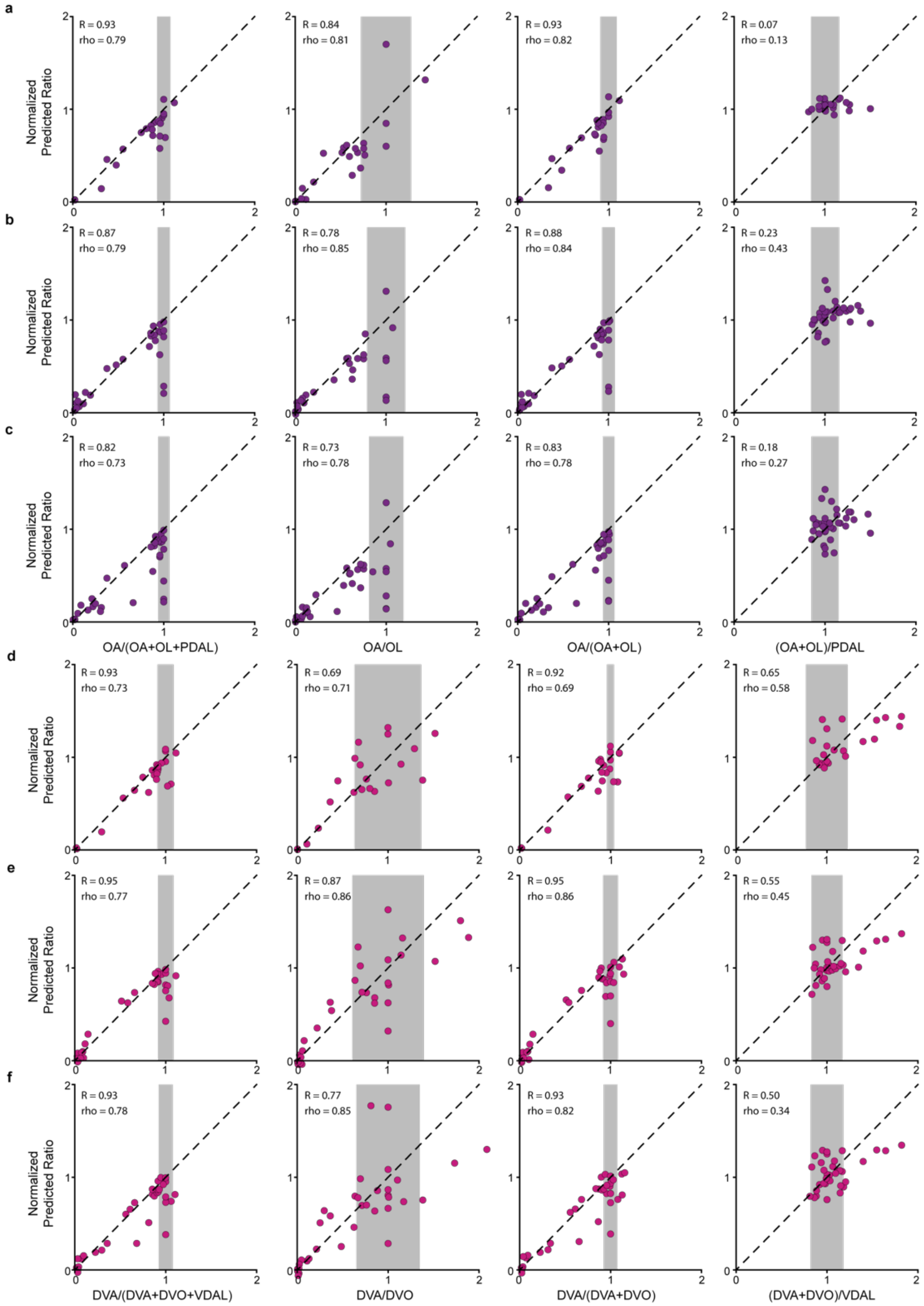


Supplementary Figure 10: Multi-task neural network predictions correlate with experimentally measured activities for CsOAC variants withheld from training. Scatter plots comparing predicted and experimentally measured values for CsOAC sequence variants withheld during supervised multi-task neural networks training during round 1 (a,d), round 2 (b,e), and round 3 (c,f). Each panel corresponds to one predicted functional output from the multi-task neural network, with variants colored by product class (OA-related outputs in purple; DVA-related outputs in pink). Points represent individual CsOAC variants. Gray shaded regions indicate the standard deviation of CsOAC observed in experiments.

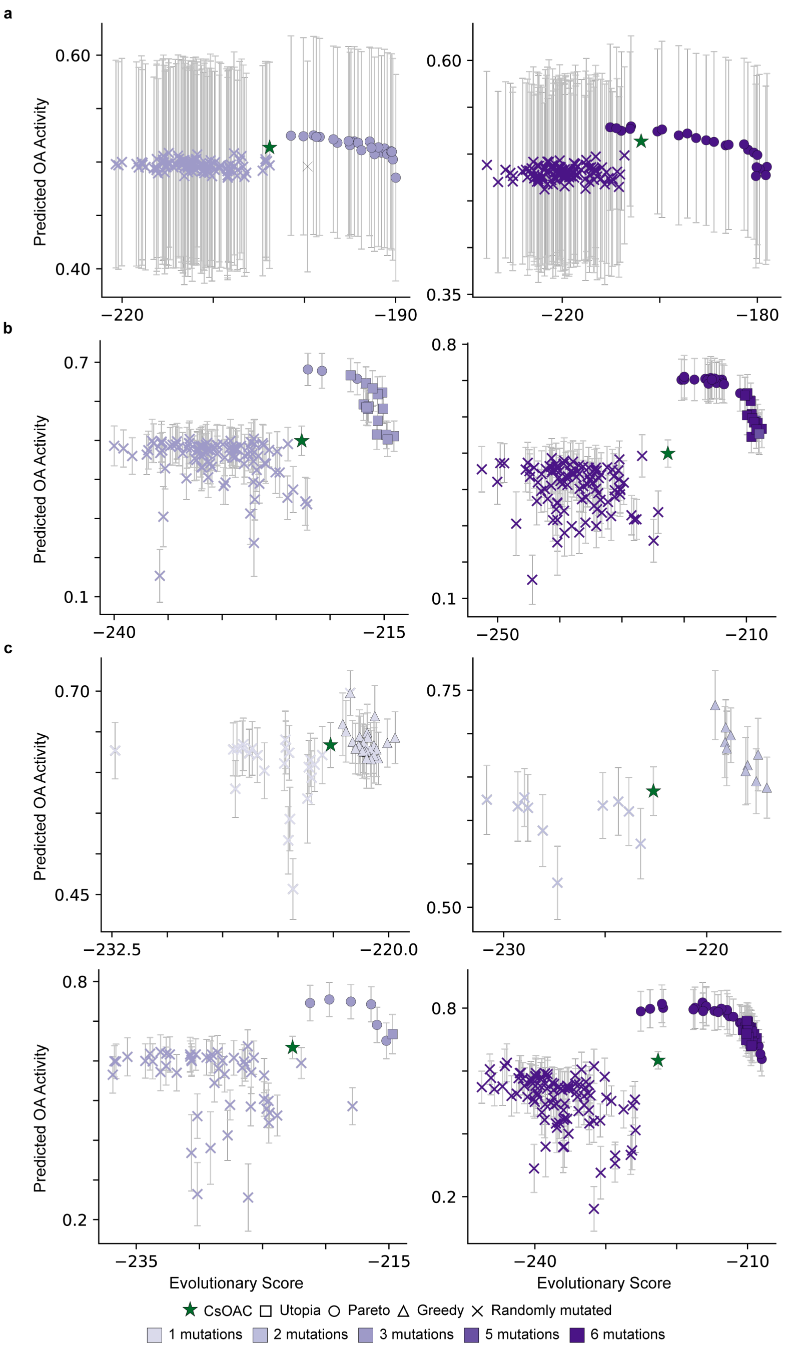


Supplementary Figure 11: Plots of multi-task neural network predicted functional scores and variational autoencoder (VAE) evolutionary scores for generated sequence designs from round 1 (a), round 2 (b), and round (3) for OA-related objectives (Supplementary Table 4-5). Each point represents a sequence variant and error bars denote standard deviation between predictions for the ensemble of multi-task neural networks. The green star denotes wild-type CsOAC. Optimization method is indicated for utopia (square), pareto (circle), greedy (triangle) with shapes. For details regarding optimization strategies, see Supplementary Table 4-5.

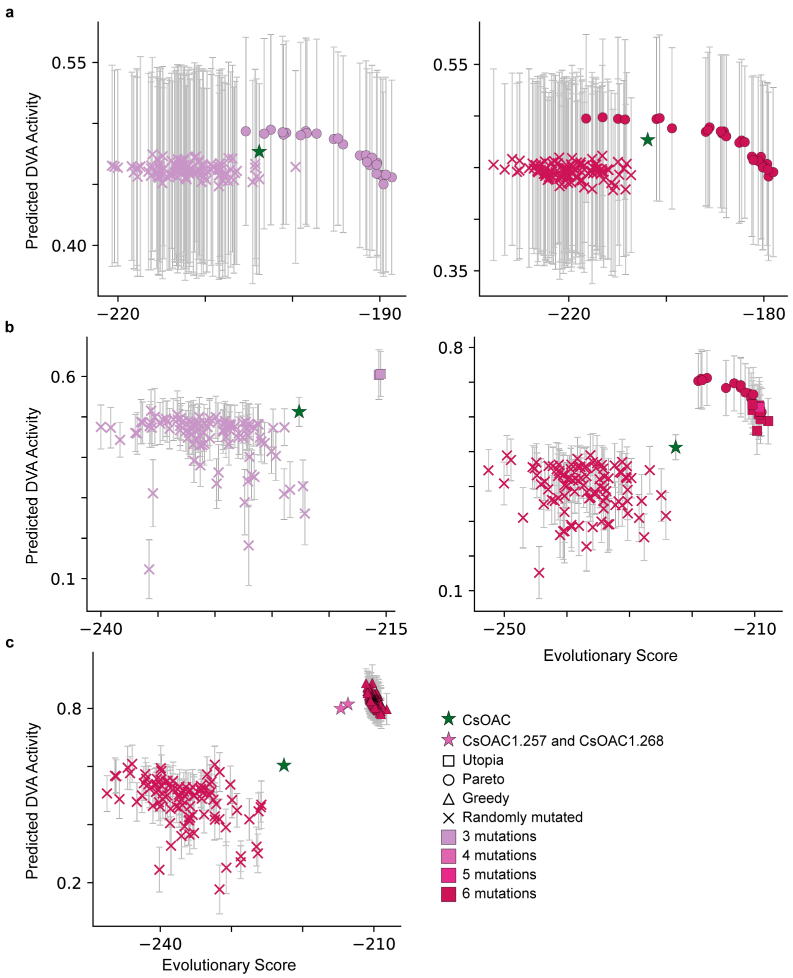


Supplementary Figure 12: Plots of multi-task neural network predicted functional scores and variational autoencoder evolutionary scores for generated sequence designs from round 1 (a), round 2 (b), and round 3 (c) for DVA-related objectives (Supplementary Table 4-5). Each point represents a sequence variant and error bars denote standard deviation between predictions for the ensemble of multi-task neural networks. The green star denotes wild-type CsOAC. Optimization method is indicated for utopia (square), pareto (circle), greedy (triangle) with shapes. For details regarding optimization strategies, see Supplementary Table 4-5.

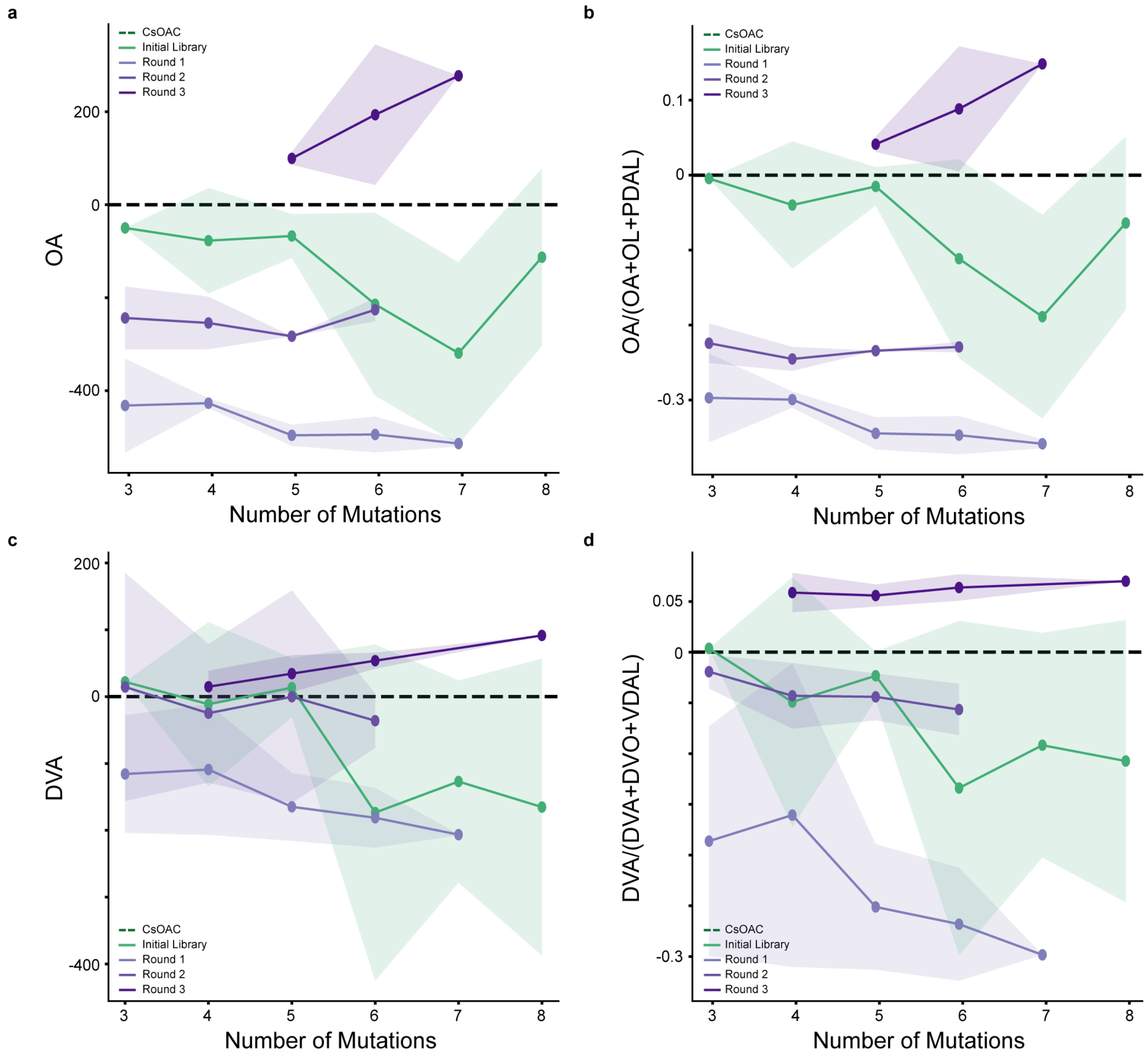


Supplementary Figure 13: Experiment Scores for Sequence Designs during ML-Driven Evolution of CsOAC Activity. a, Average titers of olivetolic acid (OA) relative to CsOAC for each library are separated by the number of mutations relative to CsOAC. b, Average OA selectivity or OA/(OA+OL+PDAL) relative to CsOAC for each library are separated by the number of mutations relative to CsOAC. c, Average titers of divarnic acid (DVA) relative to CsOAC for each library are separated by the number of mutations relative to CsOAC. d, Average DVA selectivity or DVA/(DVA+DVO+VDAL) relative to CsOAC for each library are separated by the number of mutations relative to CsOAC.

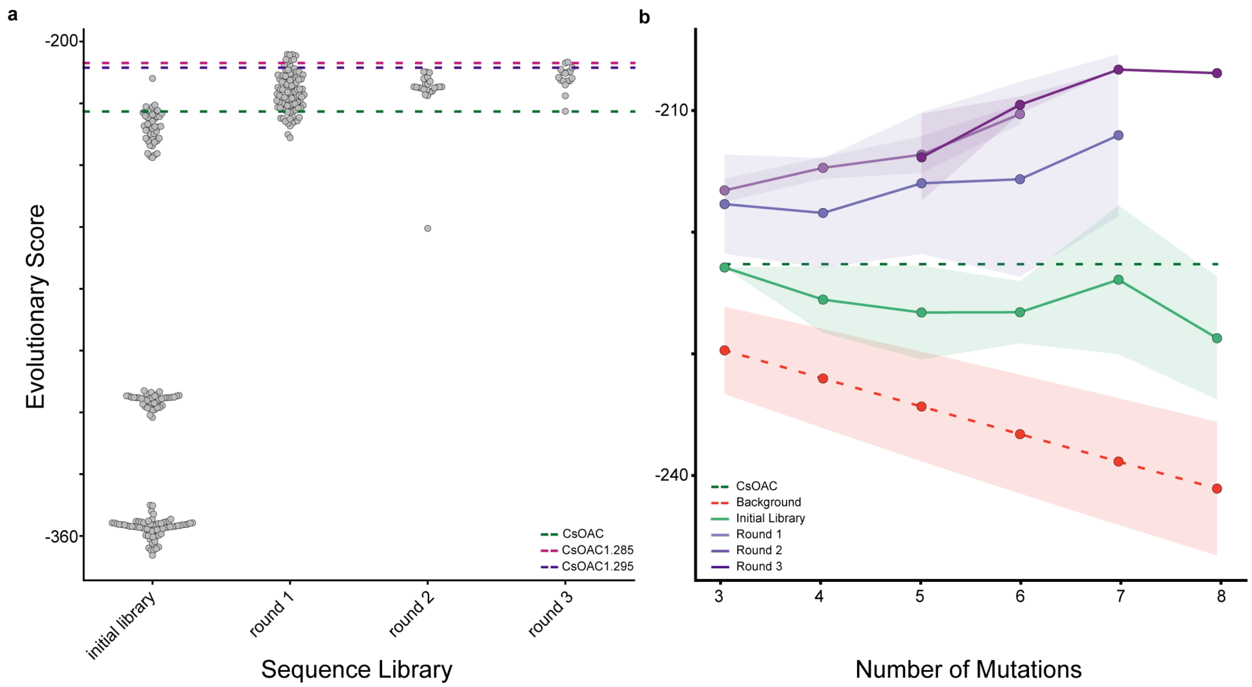


Supplementary Figure 14: Evolutionary Scores for Sequence Designs during ML-Driven Evolution of CsOAC Activity. a, Evolutionary scores for sequences designs in the initial library, round 1 sequence designs, round 2 sequence designs, and round 3 sequence designs are shown with grey points. CsOAC (green), the best sequence variant for DVA-related production CsOAC1.285 (pink), the best sequence variant for OA-related production CsOAC1.295 (purple) are indicated as baselines. b, Average evolutionary score for each library are separated by the number of mutations relative to CsOAC. Background includes sequence variants containing increasing numbers of mutations to CsOAC (Supplementary Figure 6b). Notably, this line decreases linearly with the number of mutations to CsOAC. Only sequence designs co-optimized for the VAE evolutionary score and multi-task neural network functional scores have positive slopes, suggesting a more effective exploration of the CsOAC sequence space, a challenging task given 0.2%, 0.1%, and 0.0% sequences with 6, 7, or 8 mutations (n = 500,000 each) had evolutionary scores worse than CsOAC (Supplementary Figure 6c).

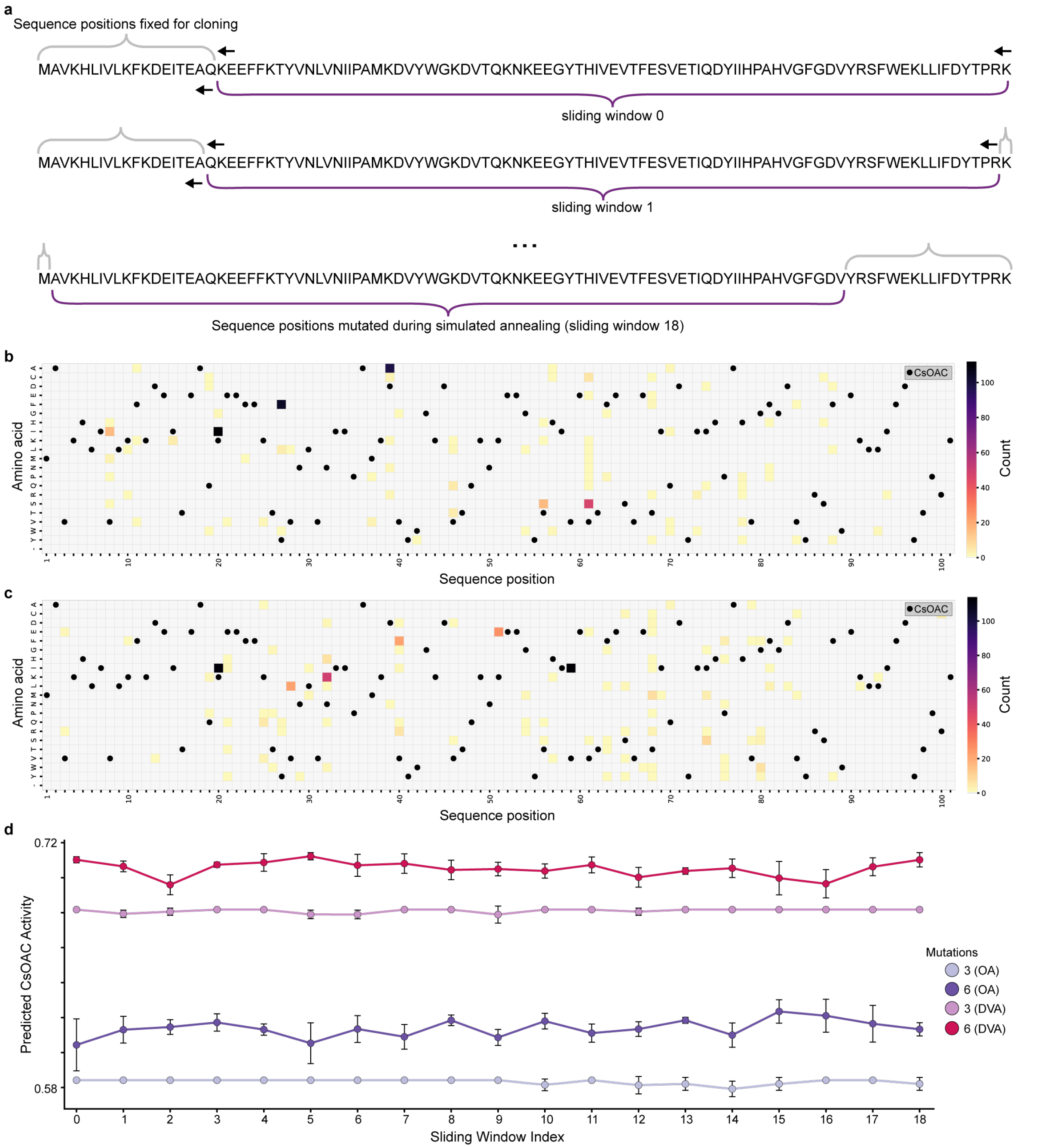


Supplementary Figure 15: Comparing sequence variants generated with different termini positions fixed for cloning. a, When generating designs with simulated annealing, we fixed the start codon and 18 additional termini positions of the CsOAC sequence. We mutated the remaining 82 sequence positions. For each sliding window, we generated sequence variants with 3 or 6 mutations to the CsOAC sequence for the functional objective OA/(OA+OL) or DVA/(DVA+DVO) via simulated annealing. b, Heatmap of sequence identity for sequence variants generated for OA/(OA+OL). c, Heatmap of sequence identity for sequence variants generated for DVA/(DVA+DVO). We selected sliding window 6 given very few mutations were observed in the fixed termini positions for both functional objectives. d, Predicted CsOAC Activity (OA/(OA+OL) or DVA/(DVA+DVO) for sequence variants with 3 or 6 mutations (n=3) for each sliding window.

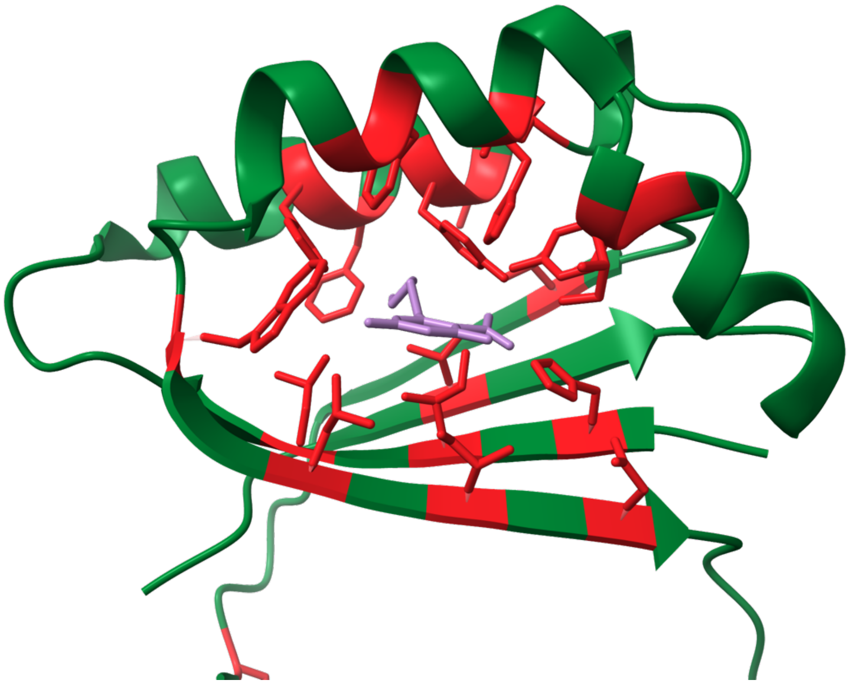

Supplementary Figure 16: Positions of the CsOAC sequence with size constraint. A size constraint applied to active-site residues defined as residues within 4 angstroms of docked OA and in the active site for sequence positions 5, 7, 9, 23-24, 27-28, 30, 40, 49, 59, 72-73, 78, 81-82, 89, 92, 94, 96 (indicated in red), where substitutions were limited to amino acids with molecular weight </= the wild-type residue size to prevent the reduction of active-site pocket volume for OA.

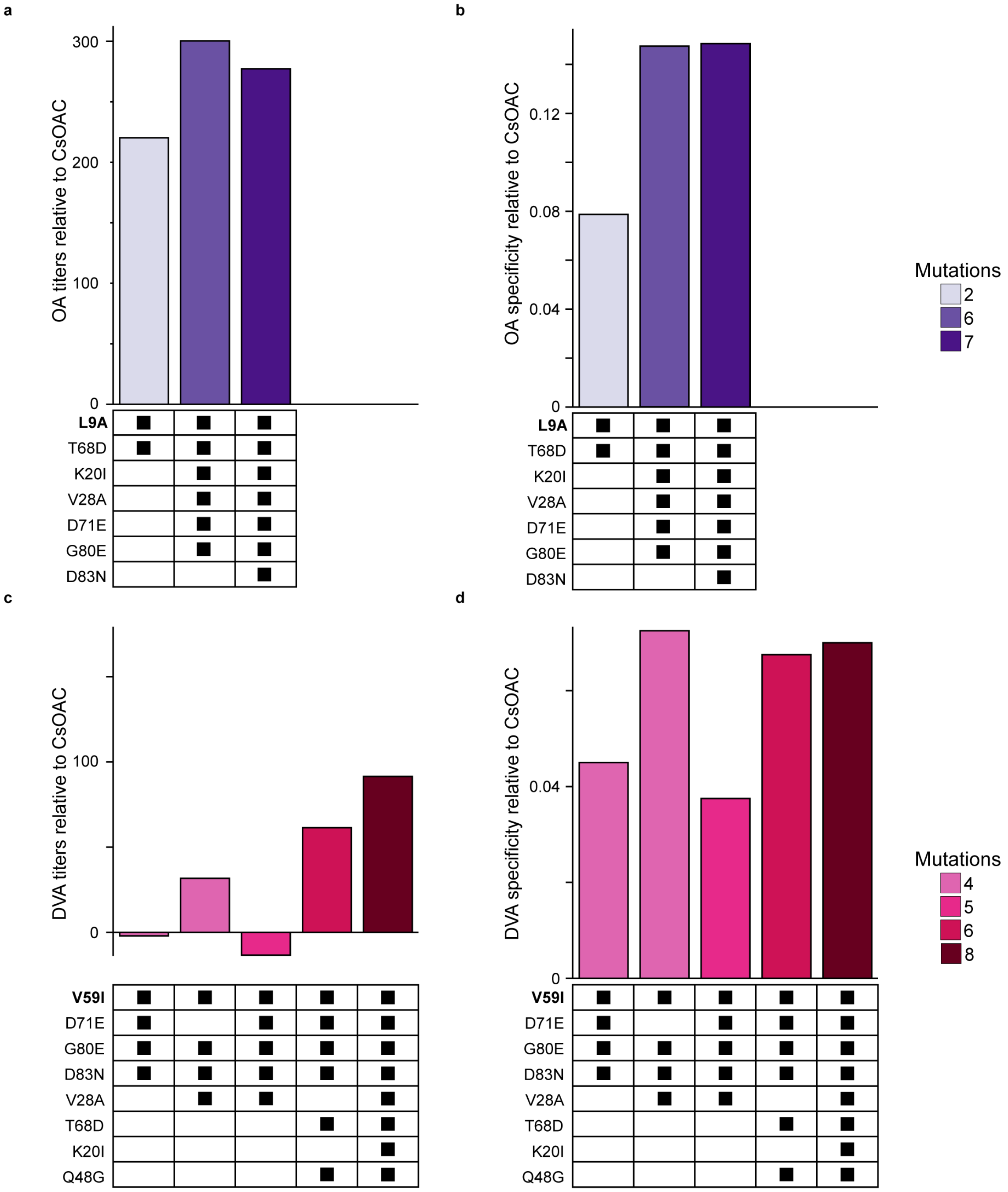


Supplementary Figure 17: Analysis of Mutational Effects from Sequence Designs in Round 3. Experimental characterization of sequence variants from round 3 containing all or a subset of L9A, T68D, K20I, V28A, D71E, G80E, and D82N for (a) olivetolic acid titers and (b) olivetolic acid specificity OA/(OA+OL+PDAL). Experimental characterization of sequence variants from round 3 containing all or a subset of V59I, D71E, G80E, D83N, V28A, T68D, K20I, and Q48G for (c) divarnic acid titers and (d) divarnic acid specificity DVA/(DVA+DVO+VDAL). The unique mutations L9A and V59I are bolded. These mutations are found in the active site and contribute to substrate selectivity.

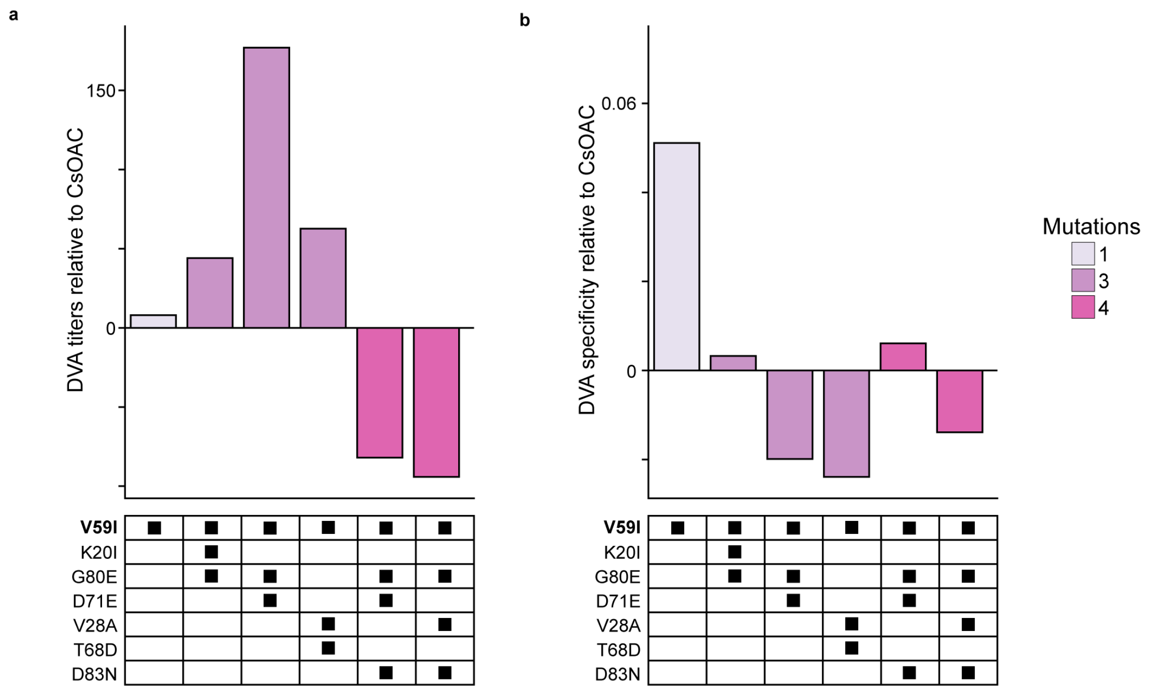


Supplementary Figure 18: Analysis of Mutational Effects from Sequence Designs in Round 2. Experimental characterization of sequence variants from round 2 containing all or a subset of V59I, D71E, G80E, D83N, V28A, T68D, K20I, and Q48G for (a) divarnic acid titers and (b) divarnic acid specificity DVA/(DVA+DVO+VDAL). The unique mutations L9A and V59I are bolded. These mutations are found in the active site and contribute to substrate selectivity.

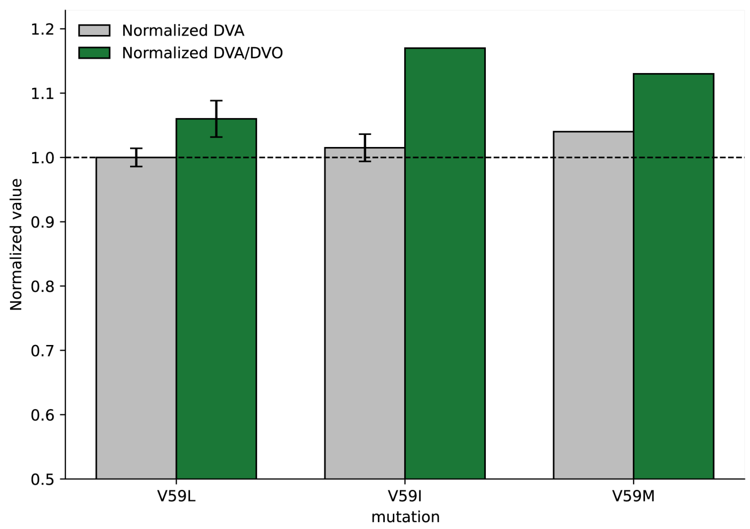


Supplementary Figure 19: Effect mutations to CsOAC V59 to leucine (L), isoleucine (I), and methionine (M). Normalized DVA titer and DVA/DVO ratio are shown for CsOAC variants containing V59L, V59I, or V59M mutations. Values are normalized relative to the CsOAC, with the dashed horizontal line indicating no change relative to the reference condition.

Supplementary Table 1: Heterologous enzymes used for cannabinoid pathway

| Enzyme/Organism/UniProt | Purpose | Source |
|---|---|---|
| **HCS2**<br>*Arabidopsis thaliana*<br>B9DGD6 | Activates hexanoic and butyric acid (Replaces CsAAE1) | Sofeo et al., 2019 introduced 3 mutations to increase activation of hexanoate |
| **PKS1.1**<br>*Cannabis sativa*<br>B1Q2B6 | Polyketide synthase (Tetraketide synthase) | We introduced 1 mutation to improve OA and DVA titers (Supplementary Figure 1). |
| **APT73.248**<br>*Streptomyces longwoodensis*<br>A0A101QTA6 | Prenylation of OA and DVA | WO 2024/137710 A3 introduced 9 mutations and a C-terminal truncation to enhance CBGA and CBGVA production |
| **THCAS***<br>*Cannabis sativa*<br>Q8GTB6 | THCA and THCVA synthesis (Replaces CsTHCAS) | WO 2023/064639 A1 replaced *C. sativa* signal peptide with signal peptide from *Y. lipolytica* (and added a C-terminal His-tag) to enhance expression |
| **ACS1**<br>*Yarrowia lipolytica*<br>Q6C2Q5 | Increases acetyl-CoA pool | WO 2024/138205 A2 increased acetyl-CoA generation with overexpression (and beta-oxidations disruptions) |
| **HMGR**<br>*Yarrowia lipolytica*<br>Q6C704 | GPP supply from the mevalonate pathway | Matthaus et al., 2014 and Cao et al., 2016 enhanced GPP production with overexpression |
| **ERG20.F88W.N119W**<br>*Yarrowia lipolytica*<br>Q8NNM1 | GPP supply from the mevalonate pathway | Ignea et al., 2013 introduced 2 mutations to increase ratio of GPP to FPP activity |
| **ERG20.A28**<br>*Yarrowia lipolytica*<br>Q8NNM1 | GPP supply from the mevalonate pathway | WO 2023/064640 A2 increased ratio of GPP to FPP activity (when Erg20 was disrupted) |
| **CNE1**<br>*Yarrowia lipolytica*<br>Q6CET3 | Unfolded THCAS response | WO 2023/064639 A1 increased THCA production with overexpression |

Supplementary Table 2: Sequence identity of heterologous enzymes

AtHCS2:

MASEENDLVFPSKEFSGQALVSSPQQYMEMHKRSMDDPAAFWSDIASEFYWKQKWGDQVFSENLDVR
KGPISIEWFKGGITNICYNCLDKNVEAGLGDKTAIHWEGNELGVDASLTYSELLQRVCQLANYLKDNGVKK
GDAVVIYLPMLMELPIAMLACARIGAVHSVVFAGFSADSLAQRIVDCKPNVILTCNAVKRGPKTINLKAIVDA
ALDQSSKDGVSVGICLTYDNSLATTRENTKWQNGRDVWWQDVISQYPTSCEVEWVDAEDPLFLLYTSG
STGKPKGVLHTTGGYMIYTATTFKYAFDYKSTDVYWCTADCGWIGGHSYVTYGPMLNGATVVVFEGAPN
YPDPGRCWDIVDKYKVSIFYTAPTLVRSLMRDDDKFVTRHSRKSLRVLGSAGEPINPSAWRWFFNVVGD
SRCPISDTWGQTETGGFMITPLPGAWPQKPGSATFPFFGVQPVIVDEKGNEIEGECSGYLCVKGSWPGA
FRTLFGDHERYETTYFKPFAGYYFSGDGCSRDKDGYYWLTGRVDDVINVSGHRIGTAEVESALVLHPQC
AEAAVVGIEHEVKGQGIYAFVTLLEGVPYSEELRKSLVLMVRNQIGAFAAPDRIHWAPGLPKTRSGKIMRR
ILRKIASRQLEELGDTSTLADPSVVDQLIALADV

YlCNE1:

MRLSKLAVSSVLAAVACAQDADAAADAAPSSQVEHPEFTPYTGAVTGFFEQFLDGHKWQKSSAMKDDE
FSYVGEWAVEEPYVFPGFKGDKGLVVKSPAAHHAITTAFDTPINNKGKTLVVQYEVKLQKGLECGGAYVK
LLSAEVNADDKGVEEFSSETPYQIMFGPDKCGSTNKVHFIVKRPLPDGTYEEKHLVSPAHARLNKLTNLY
TLVIRPKNEFEIRINGNVVKTGNLLEEGLFKPSFNPPAEIDDPEDTKPADWVEEPYMPDPEQAEKPADWD
EKAPFYIADPEAVMPADWQEDTPDYIVDPEAFKPEDWDDEEDGEWVAPEIPNPVCEEIGCGPWVAPKIQ
NPDYKGVWSQPMIENPDYKGTWAPKKIPNPNFKADEHASDLEPIGGLGFELWTMQEDILFDNIYVGHSV
DEAEAIGNATFVPKLALEAEEEKLSGPQKETAPWDTDEGVLSSVDMFLADPVSFVLERVLGFFEVFSQDP
VSAIREDPVVAAASFGLLLITSATAFGLLNVIIFLLFGKKKQSAAPAKKTKKTGDGPSKLTKADVVEAEAEAE
QAAETVVASGVDDGSAELKKRKA

CsTHCAS*:

MKFSAVSIAAALASLVAANPRENFLKCFSKHIPNNVANPKLVYTQHDQLYMSILNSTIQNLRFISDTTPKPL
VIVTPSNNSHIQATILCSKKVGLQIRTRSGGHDAEGMSYISQVPFVVVDLRNMHSIKIDVHSQTAWVEAGA
TLGEVYYWINEKNENLSFPGGYCPTVGVGGHFSGGGYGALMRNYGLAADNIIDAHLVNVDGKVLDRKSM
GEDLFWAIRGGGGENFGIIAAWKIKLVAVPSKSTIFSVKKNMEIHGLVKLFNKWQNIAYKYDKDLVLMTHFI
TKNITDNHGKNKTTVHGYFSSIFHGGVDSLVDLMNKSFPELGIKKTDCKEFSWIDTTIFYSGVVNFNTANF
KKEILLDRSAGKKTAFSIKLDYVKKPIPETAMVKILEKLYEEDVGAGMYVLYPYGGIMEEISESAIPFPHRAG
IMYELWYTASWEKQEDNEKHINWVRSVYNFTTPYVSQNPRLAYLNYRDLDLGKTNHASPNNYTQARIWG
EKYFGKNFNRLVKVKTKVDPNNFFRNEQSIPPLPPHHHHHH

YlErg20.F88W.N119W:

MSKAKFESVFPRISEELVQLLRDEGLPQDAVQWFSDSLQYNCVGGKLNRGLSVVDTYQLLTGKKELDDE
EYYRLALLGWLIELLQAFWLVSDDIMDESKTRRGQPCWYLKPKVGMIAIWDAFMLESGIYILLKKHFRQEK
YYIDLVELFHDISFKTELGQLVDLLTAPEDEVDLNRFSLDKHSFIVRYKTAYYSFYLPVVLAMYVAGITNPKD
LQQAMDVLIPLGEYFQVQDDYLDNFGDPEFIGKIGTDIQDNKCSWLVNKALQKATPEQRQILEDNYGVKD
KSKELVIKKLYDDMKIEQDYLDYEEEVVGDIKKKIEQVDESRGFKKEVLNAFLAKIYKRQK

YlErg20.A28:

MSKAKFESVFPRISEELVQLLRDEGLPQDAVQWFSDSLQYNCVGGKLNRGLSVVDTYQLLTGKKELDDE
EYYRLALLGWLIELLQAFLSDDIMDESKTRRGQPCWYLKPKVGMIAINDAFMLESGIYILLKKHFRQEKYYI
DLVELFHDISFKTELGQLVDLLTAPEDEVDLNRFSLDKHSFIVRYKTAYYSFYLPVVLAMYVAGITNPKDLQ
QAMDVLIPLGEYFQVQDDYLDNFGDPEFIGKIGTDIQDNKCSWLVNKALQKATPEQRQILEDNYGVKDKS
KELVIKKLYDDMKIEQDYLDYEEEVVGDIKKKIEQVDESRGFKKEVLNAFLAKIYKRQK

YlHMGR:

MLQAAIGKIVGFAVNRPIHTVVLTSIVASTAYLAILDIAIPGFEGTQPISYYHPAAKSYDNPADWTHIAEADIP
SDAYRLAFAQIRVSDVQGGEAPTIPGAVAVSDLDHRIVMDYKQWAPWTASNEQIASENHIWKHSFKDHV

AFSWIKWFRWAYLRLSTLIQGADNFDIAVVALGYLAMHYTFFSLFRSMRKVGSHFWLASMALVSSTFAFL
LAVVASSSLGYRPSMITMSEGLPFLVVAIGFDRKVNLASEVLTSKSSQLAPMVQVITKIASKALFEYSLEVA
ALFAGAYTGVPRLSQFCFLSAWILIFDYMFLLTFYSAVLAIKFEINHIKRNRMIQDALKEDGVSAAVAEKVAD
SSPDAKLDRKSDVSLFGASGAIAVFKIFMVLGFLGLNLINLTAIPHLGKAAAAAQSVTPITLSPELLHAIPASV
PVVVTFVPSVVYEHSQLILQLEDALTTFLAACSKTIGDPVISKYIFLCLMVSTALNVYLFGATREVVRTQSVK
VVEKHVPIVIEKPSEKEEDTSSEDSIELTVGKQPKPVTETRSLDDLEAIMKAGKTKLLEDHEVVKLSLEGKL
PLYALEKQLGDNTRAVGIRRSIISQQSNTKTLETSKLPYLHYDYDRVFGACCENVIGYMPLPVGVAGPMNI
DGKNYHIPMATTEGCLVASTMRGCKAINAGGGVTTVLTQDGMTRGPCVSFPSLKRAGAAKIWLDSEEGL
KSMRKAFNSTSRFARLQSLHSTLAGNLLFIRFRTTTGDAMGMNMISKGVEHSLAVMVKEYGFPDMDIVSV
SGNYCTDKKPAAINWIEGRGKSVVAEATIPAHIVKSVLKSEVDALVELNISKNLIGSAMAGSVGGFNAHAA
NLVTAIYLATGQDPAQNVESSNCITLMSNVDGNLLISVSMPSIEVGTIGGGTILEPQGAMLEMLGVRGPHIE
TPGANAQQLARIIASGVLAAELSLCSALAAGHLVQSHMTHNRSQAPTPAKQSQADLQRLQNGSNICIRS

YlACS1:
MSEDHPAIHPPSEFKDNHPHFGGPHLDCLQDYHQLHKESIEDPKAFWKKMANELISWSTPFETVRSGGF
EHGDVAWFPEGQLNASYNCVDRHAFANPDKPAIIFEADEPGQGRIVTYGELLRQVSQVAATLRSFGVQK
GDTVAVYLPMIPEAIVTLLAITRIGAVHSVIFAGFSSGSLRDRINDAKSKVVVTTDASMRGGKTIDTKKIVDE
ALRDCPSVTHTLVFRRAGVENLAWTEGRDFWWHEEVVKHRPYLAPVPVASEDPIFLLYTSGSTGTPKGL
AHATGGYLLGAALTAKYVFDIHGDDKLFTAGDVGWITGHTYVLYGPLMLGATTVVFEGTPAYPSFSRYWD
IVDDHKITHFYVAPTALRLLKRAGTHHIKHDLSSLRTLGSVGEPIAPDVWQWYNDNIGRGKAHICDTYWQT
ETGSHIIAPMAGVTPTKPGSASLPVFGIDPVIIDPVSGEELKGNNVEGVLALRSPWPSMARTVWNTHERY
METYLRPYPGYYFTGDGAARDNDGFYWIRGRVDDVVNVSGHRLSTAEIEAALIEHAQVSESAVVGVHDD
LTGQAVNAFVALKNPVEDVDALRKELVVQVRKTIGPFAAPKNVIIVDDLPKTRSGKIMRRILRKVLAGEEDQ
LGDISTLANPDVVQTIIEVVHSLKK

CsPKS1.1:
MNHLRAEGPASVLAIGTANPENILLQDEFPDYYFRVTKSEHMTQLKEKFRKICDKSMIRKRNCFLNEEHLK
QNPRLVEHEMQTLDARQDMLVVEVPKLGKDACAKAIKEWGQPKSKITHLIFTSASTTDMPGADYHCAKLL
GLSPSVKRVMMYQLGCYGGGTVLRIAKDIAENNKGARVLAVCCDIMACLFRGPSESDLELLVGQAIFGDG
AAAVIVGAEPDESVGERPIFELVSTGQTILPNSEGTIGGHIREAGLIFDLHKDVPMLISNNIEKCLIEAFTPIGI
SDWNSIFWITHPGGKAILDKVEEKLHLKSDKFVDSRHVLSEHGNMSSSCVLFVMDELRKRSLEEGKSTTG
DGFEWGVLFGFGPGLTVERVVVRSVPIKY

SlAPT73.248:
MDEVYAAVEQTSRLLDVPCSPDRFEPVWKAFGDQLPDSHLLFSMAAGEAHRGELDFDFSLRPEGADPY
TTALEHGFIEPTDHPVGSVLAEVGKRFAIASYGVEYGVVGGFKKSYAFFPLDDFPPLAEFARIPSVPPCLA
GHVETLTRLGFDDKVSIIGVNYQKKTLNVYLAASAVDTGDKLALLRAFGYPEPDARVRQFIPREFVSGVAL
APSGASYYKLAAIYQKGRRPLDA

CsOAC:
MAVKHLIVLKFKDEITEAQKEEFFKTYVNLVNIIPAMKDVYWGKDVTQKNKEEGYTHIVEVTFESVETIQDYI
IHPAHVGFGDVYRSFWEKLLIFDYTPRK

CsOAC1.285:
MAVKHLIVLKFKDEITEAQIEEFFKTYANLVNIIPAMKDVYWGKDVTGKNKEEGYTHIIEVTFESVEDIQEYII
HPAHVEFGNVYRSFWEKLLIFDYTPRK

CsOAC1.295:
MAVKHLIVAKFKDEITEAQIEEFFKTYANLVNIIPAMKDVYWGKDVTQKNKEEGYTHIVEVTFESVEDIQEYII
HPAHVEFGDVYRSFWEKLLIFDYTPRK

Supplementary Table 3: Genetic modification defining *Yarrowia lipolytica* chassis strain SB3829

| Gene | Modification | Source | Purpose |
|---|---|---|---|
| **AtHCS2** | Overexpression | Sofeo et al., 2019 | Increase activation of hexanoic and butyric acid |
| **CsTHCAS*** | Overexpression | WO 2023/064639 A1 | Increase THCA and THCVA synthesis |
| **YlACS1** | Overexpression | WO 2024/138205 A2 | Increase acetyl-CoA supply |
| **YlHMGR** | Overexpression | Matthaus et al., 2014 and Cao et al., 2016 | Increase GPP supply from the mevalonate pathway |
| **YlERG20.F88W.N119W** | Overexpression | Ignea et al., 2013 | Increase GPP supply from the mevalonate pathway |
| **YlERG20.A28** | Overexpression | WO 2023/064640 A2 | Increase GPP supply from the mevalonate pathway |
| **YlCNE1** | Overexpression | WO 2023/064639 A1 | Improve THCAS folding efficiency to increase activity |
| Δ*acd1* | Disruption | WO 2024/138205 A2 | Blocks acyl-CoA dehydrogenation to increase C4/C6 yield |
| Δ*pox3* | Disruption | WO 2024/138205 A2 | Reduces beta-oxidation to increase C4/C6 yield |
| Δ*pox5* | Disruption | WO 2024/138205 A2 | Reduces beta-oxidation to increase C4/C6 yield |
| Δ*ivd1* | Disruption | WO 2024/138205 A2 | Blocks isovaleryl-CoA dehydrogenation to increase C4 yield |
| Δ*bckd* | Disruption | WO 2024/138205 A2 | Reduces undesireable isobutyl-derived byproducts from *ivd1* disruption |

### Supplementary Table 4: Experimental conditions for screening CsOAC sequence variants

| Exp | Strain | Substrate | Feed (mM) | Substrate incubation | Fermentation time | Notes |
|---|---|---|---|---|---|---|
| 0 | SB-2036 | C4 (butyric acid) | 0+3 | 6 hr | 30 hr | Initial library |
| 1 | SB-741.PKS1.64 | C4 (butyric acid) | 0+4 | 24 hr | 48 hr | Initial library |
| 2 | SB-2036 | C6 (hexanoic acid) | 0+3 | 6 hr | 30 hr | Initial library |
| 3 | SB-741.PKS1.64 | C6 (hexanoic acid) | 0+4 | 24 hr | 48 hr | Initial library |
| 4 | SB-741 | C4 (butyric acid) | 0+3+3 | 24 hr \| 16 hr | 48 hr | Initial library |
| 5 | SB-741 | C6 (hexanoic acid) | 0+3+3 | 24 hr \| 16 hr | 48 hr | Initial library |
| 6 | SB-741 | C4 (butyric acid) | 0+4 | 24 hr | 48 hr | Round 1 designs |
| 7 | SB-741 | C6 (hexanoic acid) | 0+4 | 24 hr | 48 hr | Round 1 designs |
| 8 | SB-741 | C4 (butyric acid) | 0+4 | 24 hr | 48 hr | Round 1 validation |
| 9 | SB-741 | C6 (hexanoic acid) | 0+4 | 24 hr | 48 hr | Round 1 validation |
| 10 | SB-741 | C4 (butyric acid) | 0+4 | 24 hr | 48 hr | Round 2 designs |
| 11 | SB-741 | C6 (hexanoic acid) | 0+4 | 24 hr | 48 hr | Round 2 designs |
| 12 | SB-3824 | C4 (butyric acid) | 0+9 | 8 hr | 32 hr | Round 3 designs |
| 13 | SB-741 | C6 (hexanoic acid) | 0+4 | 8 hr | 32 hr | Round 3 designs |
| 14 | SB-3829 | C4 (butyric acid) | 0+5+5 | 24 hr \| 18 hr | 48 hr | Best designs |
| 15 | SB-3829 | C6 (hexanoic acid) | 0+5+5 | 24 hr \| 18 hr | 48 hr | Best designs |

Feed values indicate sequential substrate additions reported using shorthand notation. For experiments with multiple incubation times (e.g., 24 hr | 16 hr), the listed values denote the duration that substrate was present prior to quenching rather than the time of addition. For example, a notation of 0+3+3 with (24 hr | 16 hr) and a 48 hr fermentation indicates no substrate for the first 24 hr, followed by 3 mM additions at 24 hr and 32 hr, corresponding to incubation periods of 24 hr and 16 hr before quenching at 48 hr.

Supplementary Table 5: Training Multi-task Neural Networks

| **Hyperparameter** | **Values (Consistent across rounds)** |
|---|---|
| Architecture | Fully connected neural network |
| Number of models in ensemble | 100 |
| Embedding | Learned embedding with 21 tokens mapped to 16-dimensional vectors |
| Optimizer | Adam |
| Learning rate | $5 \times 10^{-6}$ |
| Weight decay | $1 \times 10^{-3}$ |
| Patience | 200 (Early stopping on Validation Loss) |
| Batch size | 32 |
| Number of hidden layers | 1 |
| Activations | ReLU after first linear layer |
| Dropout | 0.2 applied after first linear layer |

| **Hyperparameter** | **Round 1** | **Round 2** | **Round 3** |
|---|---|---|---|
| Epochs | 100 | 10,000 | 10,000 |
| Number of experiments ($n_{\text{exps}}$) | 6 | 10 | 12 |
| Number of labels ($n_{\text{labels}} = 4n_{\text{exps}}$) | 24 | 40 | 48 |
| Loss-weight pattern | [0.2, 1, 1, 0.1] | [1, 1, 1, 0.1] | [1, 1, 1, 0.1] |
| Hidden layer width | 2000 | 65 | 65 |
| Output dimension | 24 | 40 | 48 |

### Supplementary Table 6: Training Variational Autoencoder

| **Hyperparameter** | **Value** |
|---|---|
| Input | Integer-encoded amino-acid sequence of length $s_{\mathrm{len}}$ (WT length) with a 21-token alphabet. |
| Embedding | Learned embedding with 21 tokens mapped to 16-dimensional vectors. Output shape: $(B, s_{\mathrm{len}}, 16)$. |
| Encoder: permute | Swap to channels-first for convolutions. Shape: $(B, 16, s_{\mathrm{len}})$. |
| Encoder: Conv1 | 1D convolution with 16 input channels, 32 output channels, kernel size $k_s = 4$, followed by leaky ReLU. |
| Encoder: Conv2 | 1D convolution with 32 input channels, 64 output channels, kernel size $k_s = 4$, followed by leaky ReLU. |
| Encoder: flatten | Flatten to a vector of size $n_{\mathrm{param}} = (s_{\mathrm{len}} - 2(k_s - 1)) \times 64$. |
| Encoder: dense | Fully connected layer mapping $n_{\mathrm{param}} \to 1000$, followed by ReLU. |
| Latent heads | Two linear layers mapping $1000 \to n_{\mathrm{latent}}$ to produce $\mu$ and $\log \sigma^2$, with $n_{\mathrm{latent}} = 40$. |
| Reparameterization | Sample $z = \mu + \epsilon \exp\left(\frac{1}{2} \log \sigma^2\right)$ with $\epsilon \sim \mathcal{N}(0, I)$. |
| Decoder: dense 1 | Fully connected layer mapping $n_{\mathrm{latent}} \to 1000$, followed by ReLU. |
| Decoder: dense 2 | Fully connected layer mapping $1000 \to n_{\mathrm{param}}$, then reshape to $(B, 64, s_{\mathrm{len}} - 2(k_s - 1))$. |
| Decoder: Deconv1 | 1D transposed convolution with 64 input channels, 32 output channels, kernel size $k_s = 4$, followed by leaky ReLU. |
| Decoder: Deconv2 | 1D transposed convolution with 32 input channels, 16 output channels, kernel size $k_s = 4$, followed by leaky ReLU. |
| Decoder: output projection | Linear projection at each position mapping 16 features to 21 token logits (per-position categorical distribution). |
| Training hyperparameters | Batch size 64; epochs 50; optimizer Adam; learning rate $7.5 \times 10^{-6}$. |

Supplementary Table 7: Computing Functional Scores during Simulated Annealing

| Round | Objective | Targets used in score | Ensemble aggregation |
|---|---|---|---|
| 1 | DVA (C4) | `4_DVA/DVO` and `4_DVA/(DVA+DVO)` | Ensemble mean: $\mu_k = \frac{1}{100}\sum_{j=1}^{100} \hat{y}_k^{(j)}$ |
| 1 | OA (C6) | `5_OA/OL` and `5_OA/(OA+OL)` | Ensemble mean: $\mu_k = \frac{1}{100}\sum_{j=1}^{100} \hat{y}_k^{(j)}$ |
| 2 | DVA (C4) | `4_DVA/C4` and `6_DVA/C4` | LCB (5th percentile): $\mathrm{LCB}_k = Q_{0.05}\left(\{\hat{y}_k^{(j)}\}_{j=1}^{100}\right)$ |
| 2 | OA (C6) | `5_OA/C6` and `7_OA/C6` | LCB (5th percentile): $\mathrm{LCB}_k = Q_{0.05}\left(\{\hat{y}_k^{(j)}\}_{j=1}^{100}\right)$ |
| 3 | DVA (C4) | `4_DVA/(DVA+DVO)` and `6_DVA/(DVA+DVO)` | LCB (5th percentile): $\mathrm{LCB}_k = Q_{0.05}\left(\{\hat{y}_k^{(j)}\}_{j=1}^{100}\right)$ |
| 3 | OA (C6) | `7_OA/(OA+OL)` and `9_OA/(OA+OL)` | LCB (5th percentile): $\mathrm{LCB}_k = Q_{0.05}\left(\{\hat{y}_k^{(j)}\}_{j=1}^{100}\right)$ |

Outputs were selected based on (i) reliability, assessed by the number of training variants with observed measurements for each output and (ii) preference for readouts expected to increase under improved DVA or OA production.

### Supplementary Table 8: Sequence Design Parameters via Simulated Annealing

| Round | Target | Initial Sequence | $N_{\mathrm{mut}}$ | $\lambda$ | $T_0$ | $T_f$ | $n_{\mathrm{steps}}$ | Strategy | Trials | Constraints |
|---|---|---|---|---|---|---|---|---|---|---|
| 1 | DVA | CsOAC | 3 | 2 | $10^{-0.75}$ | $10^{-4}$ | 10 k | Pareto | 50 | C1 |
| 1 | OA | CsOAC | 3 | 2 | $10^{-0.75}$ | $10^{-4}$ | 10 k | Pareto | 50 | C1 |
| 1 | DVA | CsOAC | 6 | 3 | $10^{-1}$ | $10^{-4}$ | 30 k | Pareto | 35 | C1 |
| 1 | OA | CsOAC | 6 | 3 | $10^{-1}$ | $10^{-4}$ | 30 k | Pareto | 35 | C1 |
| 2 | DVA | CsOAC | 3 | 2 | $10^{-0.5}$ | $10^{-2.75}$ | 50 k | Pareto | 50 | C2 |
| 2 | OA | CsOAC | 3 | 2 | $10^{-0.5}$ | $10^{-2.75}$ | 50 k | Pareto | 50 | C2 |
| 2 | DVA | CsOAC | 6 | 3 | $10^{-0.5}$ | $10^{-2.75}$ | 100 k | Pareto | 50 | C2 |
| 2 | OA | CsOAC | 6 | 3 | $10^{-0.5}$ | $10^{-2.75}$ | 100 k | Pareto | 50 | C2 |
| 2 | DVA | CsOAC | 3 | 2 | $10^{-0.5}$ | $10^{-2.75}$ | 40 k | Utopia | 150 | C2 |
| 2 | OA | CsOAC | 3 | 2 | $10^{-0.5}$ | $10^{-2.75}$ | 40 k | Utopia | 150 | C2 |
| 2 | DVA | CsOAC | 6 | 3 | $10^{-0.5}$ | $10^{-2.75}$ | 100 k | Utopia | 75 | C2 |
| 2 | OA | CsOAC | 6 | 3 | $10^{-0.5}$ | $10^{-2.75}$ | 100 k | Utopia | 75 | C2 |
| 3 | OA | CsOAC | 3 | 2 | $10^{-0.5}$ | $10^{-2.75}$ | 50 k | Pareto | 50 | C2; S |
| 3 | OA | CsOAC | 6 | 3 | $10^{-0.5}$ | $10^{-2.75}$ | 100 k | Pareto | 50 | C2; S |
| 3 | OA | CsOAC | 3 | 1 | $10^{-0.5}$ | $10^{-2.25}$ | 30 k | Utopia | 150 | C2; S |
| 3 | OA | CsOAC | 6 | 3 | $10^{-0.5}$ | $10^{-2.75}$ | 100 k | Utopia | 75 | C2; S |
| 3 | OA | CsOAC | 1 | - | - | - | - | Greedy | - | C2; S |
| 3 | OA | CsOAC | 2 | - | - | - | - | Greedy | - | C2; S |
| 3 | DVA | CsOAC | 2 | - | - | - | - | Greedy | - | C2 |
| 3 | DVA | CsOAC1.257 | 2 | - | - | - | - | Greedy | - | C2 |
| 3 | DVA | CsOAC1.268 | 2 | - | - | - | - | Greedy | - | C2 |

Design parameters used for simulated annealing–based sequence optimization across all experimental rounds. $N_{\mathrm{mut}}$ denotes the number of mutations introduced relative to the initial sequence. $\lambda$ is the mutational rate for the Poisson distribution. $T_0$ and $T_f$ denote the initial and final temperatures for the logarithmic temperature gradient, and $n_{\mathrm{steps}}$ indicates the total number of iterations per trial (simulation). Pareto optimization performs weighted multi-objective sweeps of tradeoff coefficients. Utopia optimization minimizes Euclidean distance to the ideal objective point. Greedy denotes exhaustive enumeration of all allowed mutants for a given $N_{\mathrm{mut}}$. Trials correspond to independent optimization runs per parameter configuration (e.g., distinct Pareto weightings or repeated utopia searches). C1 refers to the cloning constraint that prevented mutations to sequence positions 1-16 at the N-terminus. C2 refers to the cloning constraint that prevented mutations to sequence positions 1-7 at the N-terminus (MAVKHLI–) and 89-101 at the C-terminus (–EKLLIFDYTPRK). S refers to the size constraint applied to active-site residues defined as residues within 4 angstroms of docked OA and in active site for sequence positions 5, 7, 9, 23-24, 27-28, 30, 40, 49, 59, 72-73, 78, 81-82, 89, 92, 94, 96, where substitutions were limited to amino acids with molecular weight $\leq$ the wild-type residue to prevent the reduction of active-site pocket volume for OA.

Supplementary Table 9: Related cannabinoid production reported in literature

| **Author** | **Year** | **Compound** | **Platform** | **Scale / Feed** | **Production** |
|---|---|---|---|---|---|
| Luo et al. | 2019 | THCA | *Saccharomyces cerevisiae* | 24-deep-well plates; galactose | 8 mg/L in strain yCAN53. |
| Luo et al. | 2019 | THCVA | *Saccharomyces cerevisiae* | 24-deep-well plates; galactose | 4.8 mg/L in strain yCAN53. |
| Ma et al. | 2022 | OA | *Yarrowia lipolytica* | Shaker flask; YPD (glucose) | 9.18 mg/L in strain YL128. |
| Hong et al. | 2025 | OA | *Yarrowia lipolytica* | Shaker flask; YPD (glucose) | 6.73 mg/L in strain YX104. |
| Hong et al. | 2025 | OA | *Yarrowia lipolytica* | Shaker flask; YPD (glucose) | 1.49 mg/L in strain YX109. |

1. Ignea, C., Pontini, M., Maffei, M. E., Makris, A. M. & Kampranis, S. C. Engineering Monoterpene Production in Yeast Using a Synthetic Dominant Negative Geranyl Diphosphate Synthase. *ACS Synth. Biol.* **3**, 298–306 (2014).
2. Kambourakis, S., Komor, R. S., Keul, N. D., Caiazza, N. C. & Urano, J. Optimized biosynthesis pathway for cannabinoid biosynthesis.
3. Luo, X. *et al.* Complete biosynthesis of cannabinoids and their unnatural analogues in yeast. *Nature* **567**, 123–126 (2019).
